# Population genomics-guided engineering of phenazine biosynthesis in *Pseudomonas chlororaphis*

**DOI:** 10.1101/2023.02.16.528854

**Authors:** Sarah Thorwall, Varun Trivedi, Ian Wheeldon

**Author notes:** These authors contributed equally (Sarah Thorwall, Varun Trivedi).

## Abstract

The emergence of next-generation sequencing (NGS) technologies has made it possible to not only sequence entire genomes, but also identify metabolic engineering targets across the pangenome of a microbial population. This study leverages NGS data as well as existing molecular biology and bioinformatics tools to identify and validate genomic signatures for improving phenazine biosynthesis in *Pseudomonas chlororaphis*. We sequenced a diverse collection of 34 *Pseudomonas* isolates using short- and long-read sequencing techniques and assembled whole genomes using the NGS reads. In addition, we assayed three industrially relevant phenotypes (phenazine production, biofilm formation, and growth temperature) for these isolates in two different media conditions. We then provided the whole genomes and phenazine production data to a unitig-based microbial genome-wide association study (mGWAS) tool to identify novel genomic signatures responsible for phenazine production in *P. chlororaphis*. Post-processing of the mGWAS analysis results yielded 330 significant hits influencing the biosynthesis of one or more phenazine compounds. Based on a quantitative metric (called the phenotype score), we elucidated the most influential hits for phenazine production and experimentally validated them *in vivo* in the most optimal phenazine producing strain. Two genes significantly increased phenazine-1-carboxamide (PCN) production: a histidine transporter (ProY_1), and a putative carboxypeptidase (PS__04251). A putative MarR-family transcriptional regulator decreased PCN titer when overexpressed in a high PCN producing isolate. Overall, this work seeks to demonstrate the utility of a population genomics approach as an effective strategy in enabling identification of targets for metabolic engineering of bioproduction hosts.

## Introduction

The development of next-generation sequencing and CRISPR genome editing has enabled entire microbial genomes to be sequenced and manipulated, resulting in genome-wide metabolic engineering approaches often within non-traditional hosts. With further advancements in DNA sequencing technologies it is now economically feasible for a single research group to sequence small collections of tens to hundreds of microbial isolates, sometimes even in-house with portable sequencing devices. New metabolic engineering strategies could take advantage of this increasing accessibility of microbial whole-genome sequencing data and existing bioinformatics tools to analyze this data to identify metabolic engineering targets from a collection of genomes in a “pangenome”-wide or population genomics approach.

This work seeks to identify non-intuitive genetic targets from a collection of *Pseudomonas chlororaphis* isolates to improve phenazine production as part of a population genomics approach to metabolic engineering. Phenazines are redox-active, often colorful, secondary metabolites with applications in agriculture as antifungal agents and potential applications as redox mediators in flow cell batteries and bioelectrochemical devices (Clifford et al., 2021; Hollas et al., 2018; Rabaey et al., 2005). *P. chlororaphis* is a commercially available biocontrol strain which would make a good potential phenazine production host, as it is non-pathogenic to humans and plants, can utilize the inexpensive carbon source glycerol, and natively produces multiple phenazine derivatives. Even so, many strains have traits which are detrimental to industrial bioprocessing, such as biofilm formation and low growth temperatures. Here, we sequenced the genomes of 34 *Pseudomonas* isolates, characterized their bioprocess-relevant phenotypes (phenazine production, biofilm formation, and growth temperature), and conducted microbial genome-wide association studies (mGWAS) to select the optimal host strain for phenazine production and identify genetic manipulations that increase phenazine biosynthesis.

*P. chlororaphis* has already been successfully engineered for phenazine production, with metabolic engineering works pursuing rational design strategies. Replacing genes within the phenazine biosynthesis operon can modulate final phenazine composition and allow non-native phenazines to be produced, including 1-hydroxyphenazine (Wan et al., 2021) and phenazine-1,6-dicarboxylic acid derivatives iodinin and 1,6-dimethoxyphenazine (Guo et al., 2022)). Regulation of the phenazine biosynthesis operon provides opportunities to improve phenazine production, including *phzR* and *phzI* which directly regulate expression through quorum sensing (Yu et al., 2018) and components of the Gac/Rsm pathway (e.g. *rpeA, rsmE, lon* protease, *psrA, parS, gacA* (Li et al., 2020; Liu et al., 2016; Wan et al., 2021)) which indirectly interact with PhzR/PhzI in response to other environmental factors. Increasing carbon flux through the shikimate pathway, such as by overexpressing *aroB, aroD, aroE, ppsA*, and *tktA* (Li et al., 2020; Liu et al., 2016), also improves phenazine production by increasing flux through phenazine biosynthesis.

Our population genomics strategy uses mGWAS to identify metabolic engineering targets from our genomic and phenotypic data. GWAS correlate genomic and phenotypic datasets to identify causal genetic variants (Lees and Bentley, 2016). While GWAS are most commonly used to identify human disease risk factors, recent bioinformatics tools have been developed to adapt these studies to bacteria (Brynildsrud et al., 2016; Jaillard et al., 2018; Lees et al., 2016). We input our sequenced genomes and phenotypic data into DBGWAS (Jaillard et al., 2018), a unitig-based mGWAS tool, to identify genomic loci that are significantly associated with phenazine production. This approach requires no prior knowledge of relevant biosynthetic pathways and could identify previously unknown targets for metabolic engineering throughout a single genome and the genomes of a population of isolates. We further sought to experimentally validate the top mGWAS hits by overexpressing associated genes and measuring phenazine production with respect to the wildtype control.

## Results and Discussion

### Curating a *P. chlororaphis* strain collection

We purchased all unique strains of *P. chlororaphis* that were accessible to us from international culture collections, resulting in 26 strains purchased from three culture collections (**Table S1**). Eight of these strains had multiple colony morphologies present which appeared to vary in pigment production. Because these variants could differ in the phenotypes of interest and consequently may have associated genetic variation, each of these variants was treated as a separate isolate with the strain number indicating the original culture collection strain designation followed by a superscript ^1^ and ^2^ arbitrarily assigned to the differing morphologies. 16s rRNA sequencing identified 33 isolates as *P. chlororaphis* and one as *P. synxantha* for a total of 34 *Pseudomonas* isolates used in this study.

### Phenotyping for phenazine production, biofilm formation, and growth temperature

For the phenazine biosynthesis phenotyping, we quantify the four phenazine compounds naturally produced by *P. chlororaphis*: 2-hydroxyphenazine (2-HP), 2-hydroxyphenazine-1-carboxylic acid (2-HPCA), phenazine-1-carboxylic acid (PCA) and phenazine-1-carboxamide (PCN) (**Fig. 1**) (Mavrodi et al., 2006). In phenazine-producing pseudomonads, the core phenazine biosynthesis operon is responsible for synthesizing PCA which serves as the precursor for other phenazine derivatives. *P. chlororaphis* strains typically produce either PCN or a combination of 2-HPCA and 2-HP depending on whether *phzH* or *phzO*, respectively, is present and functional.

**Fig. 1.**
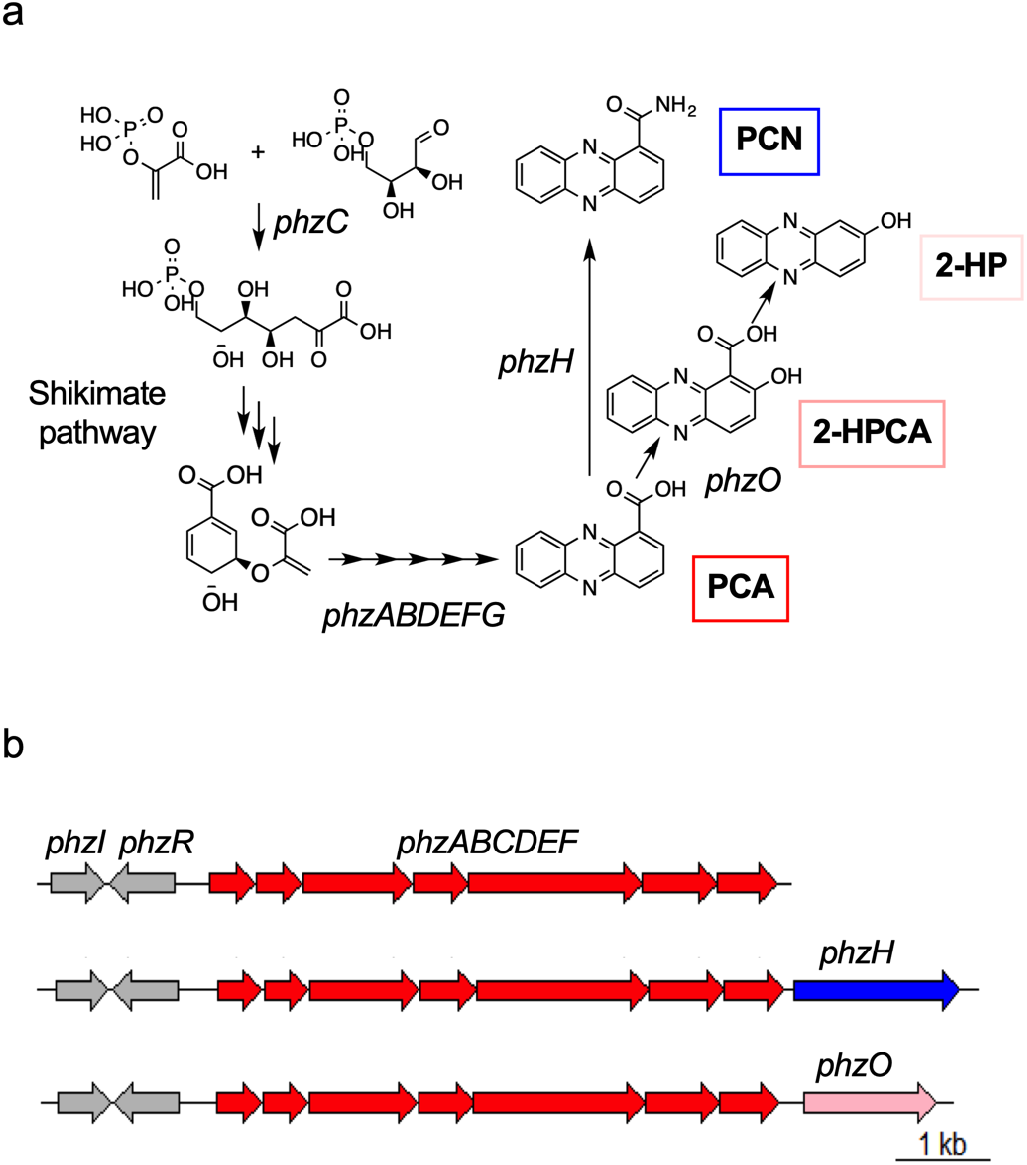
Phenazine biosynthesis pathway and operon in *P. chlororaphis*. a) Phenazine biosynthesis pathway. *P. chlororaphis* naturally produces 4 phenazines: 2-hydroxyphenazine (2-HP), 2-hydroxyphenazine-1-carboxylic acid (2-HPCA), phenazine-1-carboxylic acid (PCA) and phenazine-1-carboxamide (PCN). Chorismate from the shikimate pathway is converted into PCA. PCA can be converted into PCN or 2-HPCA by PhzH or PhzO, respectively. 2-HP is a byproduct of the spontaneous decarboxylation of 2-HPCA. b) In fluorescent pseudomonads, phenazines are produced by a highly conserved core phenazine biosynthesis operon. *phzI/phzR* encodes a two-component quorum-sensing system which regulates the expression of the *phz* operon *phzABCDEFG*, which is responsible for the production of PCA. Some strains of *P. chlororaphis* contain *phzO*, the protein product of which converts PCA into 2-HPCA (which spontaneously decomposes into 2-HP), while others contain *phzH*, which converts PCA into PCN. *phzH* or *phzO* occur immediately downstream of *phzG* in the biosynthesis operon.

We first characterized phenazine production in King’s Media B (KMB), the standard culture media for fluorescent pseudomonads (**Fig. 2**). Under these conditions, fewer than half the isolates produced more than 10 mg/L of phenazines. These low titers suggest that phenotyping in KMB may underestimate the phenazine production capacity of our strain collection. To improve phenazine production and consequently the quality of this dataset for our mGWAS analysis, we supplemented KMB with ferric iron, which has been previously reported to enhance phenazine production in some strains of *P. chlororaphis* (Chin-A-Woeng et al., 1998; van Rij et al., 2004). *KMB+Fe media improved phenazine production in 24 isolates, and 10 isolates did not produce significant phenazines in either medium. Due to its positive effects for most of the strains and its neutral effects on the remaining strains, we did additional phenotyping in KMB+Fe as well as KMB*.

**Fig. 2.**
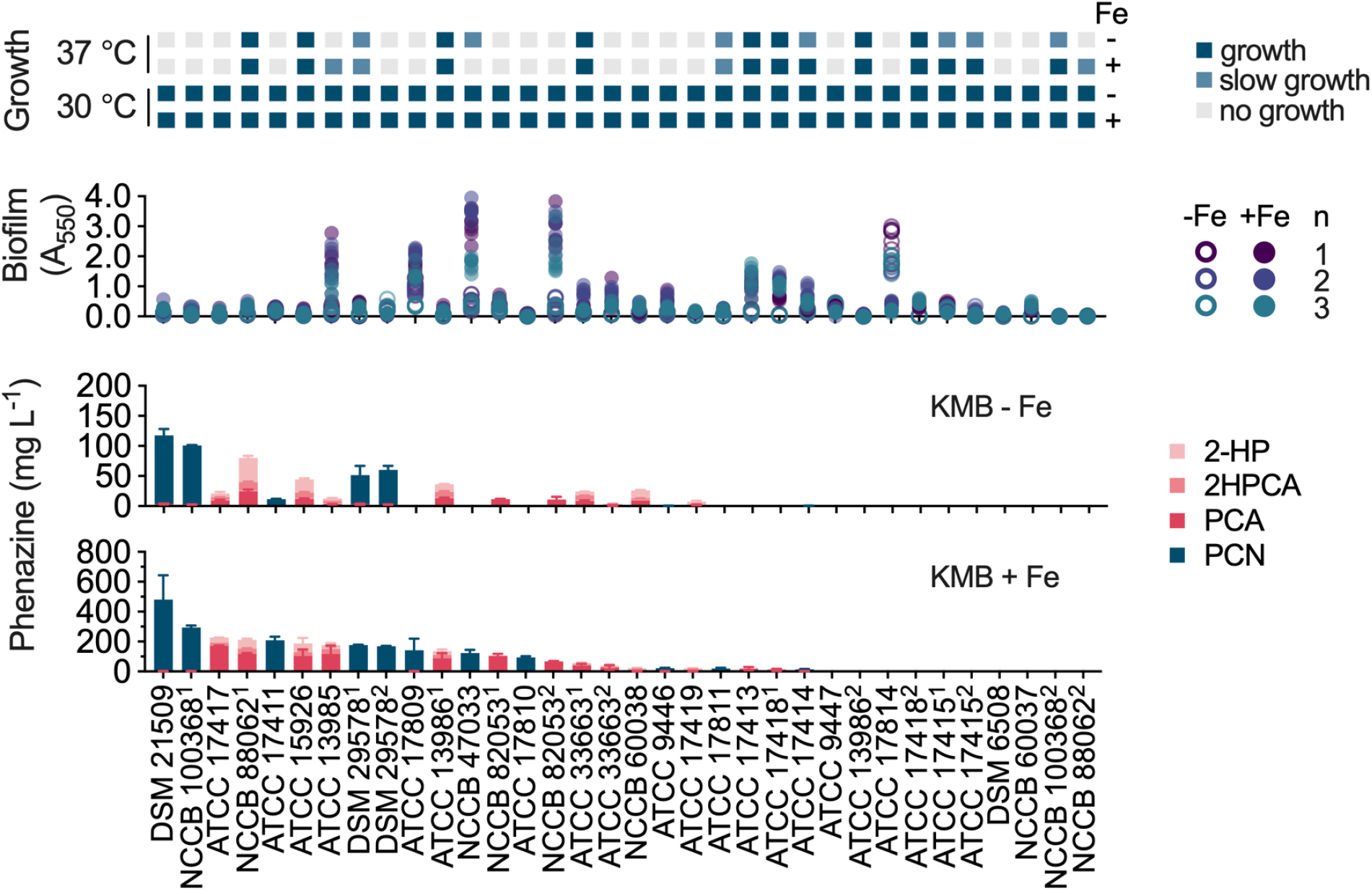
Phenazine production, biofilm formation, and growth temperature phenotyping for all isolates used in this study. All phenotyping data was collected after 48 hours of culture in either King’s Media B (-Fe) or King’s Media B + Fe (+Fe). 2-hydroxyphenazine (2-HP), 2-hydroxyphenazine-1-carboxylic acid (2-HPCA), phenazine-1-carboxylic acid (PCA) and phenazine-1-carboxamide (PCN) were quantified using HPLC. The PCN-producers primarily produced PCN, with only very small amounts of the PCA precursor detected (< 5 mg/L). The bars indicate the average of 3 replicates, and the error bars represent one standard deviation. For growth studies, dark blue indicates growth, light blue indicates slow growth, while light gray indicates no growth on solid media. For biofilm formation, each data point (A_550_, which is indicative of biofilm formation) represents a separate biological replicate, which is the average of 8 technical replicates.

Out of all strains we characterized, strain DSM 21509 was the best host strain overall for PCN and total phenazine production due to its high titers of PCN in both media conditions (477 ± 163 mg/L PCN in KMB + Fe and 114 ± 11 mg/L PCN in KMB). DSM 21509 is the type strain of *P. chlororaphis* subsp. *piscium*. This strain was isolated from the intestine of a European perch from Lake Neuchâtel, Switzerland, in 2005 (Burr et al., 2010). Strain ATCC 17417 was the top PCA (171 ± 4 mg/L) and combined PCA/2-HPCA/2-HP (∼228 mg/L) producer in KMB+Fe, while strain NCCB 88062^1^ produced the most PCA (25 ± 2 mg/L) and combined PCA/2-HPCA/2-HP in KMB. The ATCC 17417 strain was isolated from soil from a Michigan forest in 1955 (Conway et al., 1956; Stanier et al., 1966), and strain NCCB 88062 originated from the Netherlands and was deposited into NCCB in 1988 (https://wi.knaw.nl/page/NCCB_strains_display/24262). This phenazine production data for both media conditions was used as input for the mGWAS analysis.

In addition to phenazine production, we also characterized growth temperature and biofilm formation for all strains in both media conditions and used these phenotypic datasets to assess the potential of each isolate as a biotechnology host (Thorwall et al., 2020). To identify strains which could grow at common bioprocessing temperatures, we characterized growth for all strains at 30 °C and 37 °C. At 37 °C about half the strains were capable of growth, but we observed no colorful pigments (**Fig. 2**). This lack of pigments indicates that these strains produce little to no phenazines at this temperature, so the strains which can grow at 37 °C could be useful to pursue as hosts for other products but likely not for phenazines. Therefore 30 °C was selected as the fermentation temperature for this study, as all isolates could grow at the lower temperature on both media. We characterized biofilm formation to identify and rule out strains with high biofilm formation, which may be difficult to easily culture and/or clean from industrial bioreactors. In KMB, only 2 strains produced noticeable biofilm. Biofilm formation did increase in KMB+Fe, but only 5 strains had enough biofilm for it to be visibly noticeable when handling the liquid culture. Strain DSM 21509 and PCN were selected as the production strain and desired phenazine product, respectively, because this strain produced the highest overall phenazine titers, which were 99.2% PCN in KMB+Fe. Additionally DSM 21509 had low biofilm formation in both tested media conditions making it favorable to work with.

### Genome sequencing, assembly, and annotation

We sequenced all strains with both Illumina and Oxford Nanopore technologies as each technology generates reads which vary in length and accuracy, therefore affecting the quality of the resulting assemblies. Using each read set separately or together (in a hybrid approach), we assembled genomes with different assembly algorithms (*i*.*e*., SPAdes, Unicycler, Flye) to determine which algorithm and combinations of parameters yield the best assemblies. The summary statistics (*i*.*e*., number of contigs, L_50_, N_50_, assembly length, GC content, number of CDS and BUSCO score) were compared to assess genome contiguity and accuracy and thus to select the optimal genome assemblies for the mGWAS analysis (**Fig. S1**).

Contiguity statistics (number of contigs, N_50_, L_50_) describe the degree of fragmentation of an assembly. The number of contigs, or assembly fragments, should ideally approach one to accurately represent bacterial genomes with a singular circular chromosome, as is expected for *P. chlororaphis*. Genome completeness was assessed by implementing the BUSCO algorithm to calculate the percentage of expected complete and single copy orthologs which are present in each strain. The hybrid assemblies created with Unicycler were selected as the final assemblies due to their high contiguity and completeness metrics (L_50_ = 1 and BUSCO score >98% for all strains). Summary statistics for each of the final genomes, including total length, number of annotated CDS, BUSCOS scores, and assembly metrics are presented in **Table 1**. Notably, the vast majority of assemblies resulted in a single contig (22 *P. chlororaphis* and 1 *P. synxantha*), nine assembled into three or less contigs, one produced five contigs, and only one had ten contigs. Combined with the high BUSCO scores, the low number of contigs is indicative of high quality, complete genomes across our strain collection.

**Table 1.** Summary statistics for final genome assemblies. For each isolate, the assembly length, number of contigs, N_50_, L_50_, GC% and BUSCO score are reported. These assemblies were used as input to mGWAS analysis. The number of CDS was tallied from the Prokka genome annotations, and the BUSCO score was calculated as the percentage of complete and single copy BUSCOs present in each genome using the BUSCO algorithm. All other statistics were generated from QUAST. These genome assemblies are available at NCBI with the listed accession numbers. All strains are *P. chlororaphis*, except ATCC 17413 (marked with *), which is *P. synxantha*.

| Strain | Total length<br>(bp) | Contigs | N<sub>50</sub> | L<sub>50</sub> | GC (%) | CDS | BUSCOs<br>(%) | NCBI Accession<br>Number |
| --- | --- | --- | --- | --- | --- | --- | --- | --- |
| ATCC 13985 | 7 024 010 | 10 | 4 636 000 | 1 | 62.7 | 6251 | 99.1 | JAQZQZ000000000 |
| ATCC 13986<sup>1</sup> | 6 675 284 | 2 | 6 636 555 | 1 | 63.0 | 5926 | 99.5 | JAQZQY000000000 |
| ATCC 13986<sup>2</sup> | 6 682 756 | 2 | 6 644 045 | 1 | 63.0 | 5949 | 99.5 | JAQZQX000000000 |
| ATCC 15926 | 6 763 921 | 1 | 6 763 921 | 1 | 62.9 | 5998 | 99.4 | CP118156 |
| ATCC 17411 | 7 212 419 | 1 | 7 212 419 | 1 | 62.5 | 6366 | 99.2 | CP118155 |
| ATCC 17414 | 6 807 169 | 1 | 6 807 169 | 1 | 63.0 | 6048 | 99.5 | CP118154 |
| ATCC 17415<sup>1</sup> | 6 664 157 | 1 | 6 664 157 | 1 | 63.0 | 5887 | 99.4 | CP118147 |
| ATCC 17415<sup>2</sup> | 6 664 503 | 1 | 6 664 503 | 1 | 63.0 | 5884 | 99.2 | CP118146 |
| ATCC 17417 | 6 746 536 | 1 | 6 746 536 | 1 | 62.9 | 5954 | 99.1 | CP118145 |
| ATCC 17418<sup>1</sup> | 6 883 267 | 1 | 6 883 267 | 1 | 62.8 | 6075 | 99.5 | CP118144 |
| ATCC 17418<sup>2</sup> | 6 881 643 | 1 | 6 881 643 | 1 | 62.8 | 6074 | 99.5 | CP118143 |
| ATCC 17419 | 6 608 598 | 5 | 4 662 896 | 1 | 62.7 | 5919 | 99.0 | JAQZQW000000000 |
| ATCC 17809 | 7 020 903 | 1 | 7 020 903 | 1 | 62.4 | 6223 | 99.0 | CP118142 |
| ATCC 17810 | 6 863 056 | 2 | 6 791 445 | 1 | 62.7 | 6074 | 99.4 | JAQZQV000000000 |
| ATCC 17811 | 7 189 114 | 1 | 7 189 114 | 1 | 62.4 | 6422 | 99.4 | CP118153 |
| ATCC 17814 | 6 807 913 | 1 | 6 807 913 | 1 | 63.0 | 6050 | 99.4 | CP118141 |
| ATCC 33663<sup>1</sup> | 7 109 352 | 1 | 7 109 352 | 1 | 62.9 | 6281 | 99.1 | CP118152 |
| ATCC 33663<sup>2</sup> | 7 108 820 | 1 | 7 108 820 | 1 | 62.9 | 6284 | 99.2 | CP118140 |
| ATCC 9446 | 6 637 791 | 1 | 6 637 791 | 1 | 63.0 | 5909 | 99.2 | CP118151 |
| ATCC 9447 | 6 807 068 | 3 | 6 677 872 | 1 | 63.0 | 6048 | 99.4 | JAQZQU000000000 |
| DSM 21509 | 7 064 975 | 1 | 7 064 975 | 1 | 62.7 | 6246 | 99.1 | CP118150 |
| DSM 29578<sup>1</sup> | 7 216 947 | 1 | 7 216 947 | 1 | 62.5 | 6378 | 99.1 | CP118139 |
| DSM 29578<sup>2</sup> | 7 216 571 | 1 | 7 216 571 | 1 | 62.5 | 6380 | 99.1 | CP118138 |
| DSM 6508 | 7 915 166 | 3 | 7 476 725 | 1 | 62.5 | 7222 | 99.4 | JAQZQT000000000 |
| NCCB 100368<sup>1</sup> | 6 870 522 | 2 | 6 455 838 | 1 | 62.8 | 6010 | 99.2 | JAQZQS000000000 |
| NCCB 100368<sup>2</sup> | 6 870 415 | 2 | 6 455 628 | 1 | 62.8 | 6010 | 99.1 | JAQZQR000000000 |
| NCCB 47033 | 7 221 530 | 1 | 7 221 530 | 1 | 62.4 | 6378 | 99.2 | CP118137 |
| NCCB 60037 | 6 977 278 | 1 | 6 977 278 | 1 | 62.7 | 6209 | 99.1 | CP118149 |
| NCCB 60038 | 6 979 353 | 1 | 6 979 353 | 1 | 62.7 | 6214 | 99.1 | CP118136 |
| NCCB 82053<sup>1</sup> | 6 763 242 | 2 | 6 274 577 | 1 | 62.9 | 6002 | 99.4 | JAQZQQ000000000 |
| NCCB 82053<sup>2</sup> | 6 762 156 | 1 | 6 762 156 | 1 | 62.9 | 6002 | 99.4 | CP118135 |
| NCCB 88062<sup>1</sup> | 7 025 460 | 2 | 6 660 177 | 1 | 62.8 | 6233 | 99.1 | JAQZQP000000000 |
| NCCB 88062<sup>2</sup> | 6 923 225 | 1 | 6 923 225 | 1 | 62.8 | 6138 | 98.2 | CP118148 |
| *ATCC 17413 | 6 147 644 | 1 | 6 147 644 | 1 | 60.0 | 5459 | 99.9 | CP118134 |

### Assembling the *P. chlororaphis* pangenome

While the 33 *P. chlororaphis* isolates are members of the same species, their gene content varies among isolates. Comparing the assembly summary statistics (Table 1) reveals a wide range of assembly size (6.6 Mbp to 7.9 Mbp) and number of CDS (5884 to 7222 annotated CDS) so we decided to assemble the pangenome to gain further insight into these genomic differences. We input the Prokka annotations for the final *P. chlororaphis* genomes into the algorithm PEPPAN (Zhou et al. 2020) to calculate the pangenome, the total gene content of the strains. The pangenome contains 11527 total CDS and 4406 CDS common to all strains (**Fig. 3a**). This translates to 61-75% of the CDS in each genome being common to all strains, the core *P. chlororaphis* genome. The remaining CDS are members of the accessory genome (genes present in some strains and absent in others), which corresponds to the majority of this pangenome (7121 genes). These accessory genes are found in a relatively small number of strains, while all strains contain the 4406 core genome (**Fig. 3b**).

**Fig. 3.**
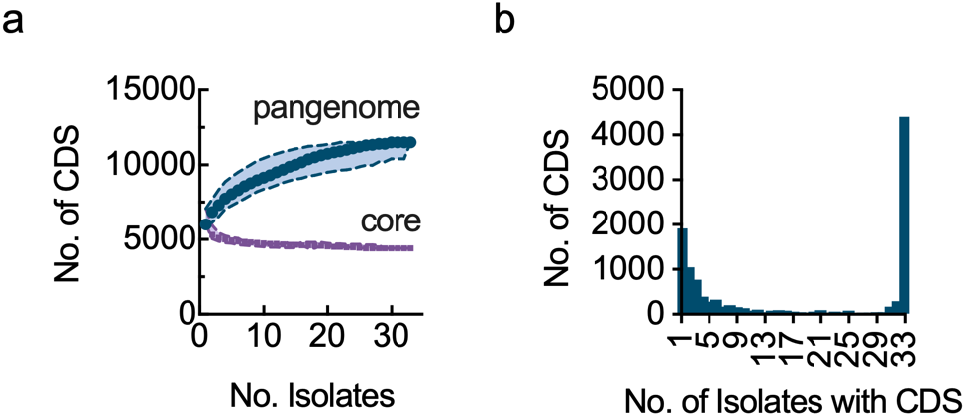
Summary of the pangenome constructed from the final hybrid genome assemblies. (a) The pangenome rarefaction curve shows how the total pangenome size (blue) increases and the core genome size (purple) decreases as isolates are added to the pangenome. (b) Histogram showing how many CDS are only found in a specific number of strains. Core genes present in almost all strains or accessory genes only found in a few are present with the highest frequencies.

In addition to the genomic variation among the strains, we observed considerable phenotypic variation as shown in Fig. 2. Most notably, strains that produced PCN did not accumulate significant quantities of PCA or other phenazine derivatives. One of the key differences amongst these strains was the presence or absence of *phzH* and *phzO*; those that produced PCN contained *phzH*, while those that produced 2-HPCA contained *phzO*, as expected (**Fig. 4**). This split amongst the population is observable in a similarity tree based on accessory gene content, which clustered the isolates containing *phzH* separately from those that contain *phzO*. In the pre-genomic era the phenotypic differences were the basis of classification; members of these two groups would likely have been classified as different *Pseudomonas* species (*e*.*g*., green pigment-producers as *P. chlororaphis*, yellow-orange pigment-producers as *P. aureofaciens* or *P. aurantiaca*; (Peix et al., 2007)). Classification in this way would incorrectly separate the groups, as our 16s and genomic sequencing shows that all 33 strains are *P. chlororaphis* with genetic variation driving the naturally produced phenazines. Collectively, the pangenome represents a large number of potential metabolic engineering targets that will be analyzed in our mGWAS.

**Fig. 4.**
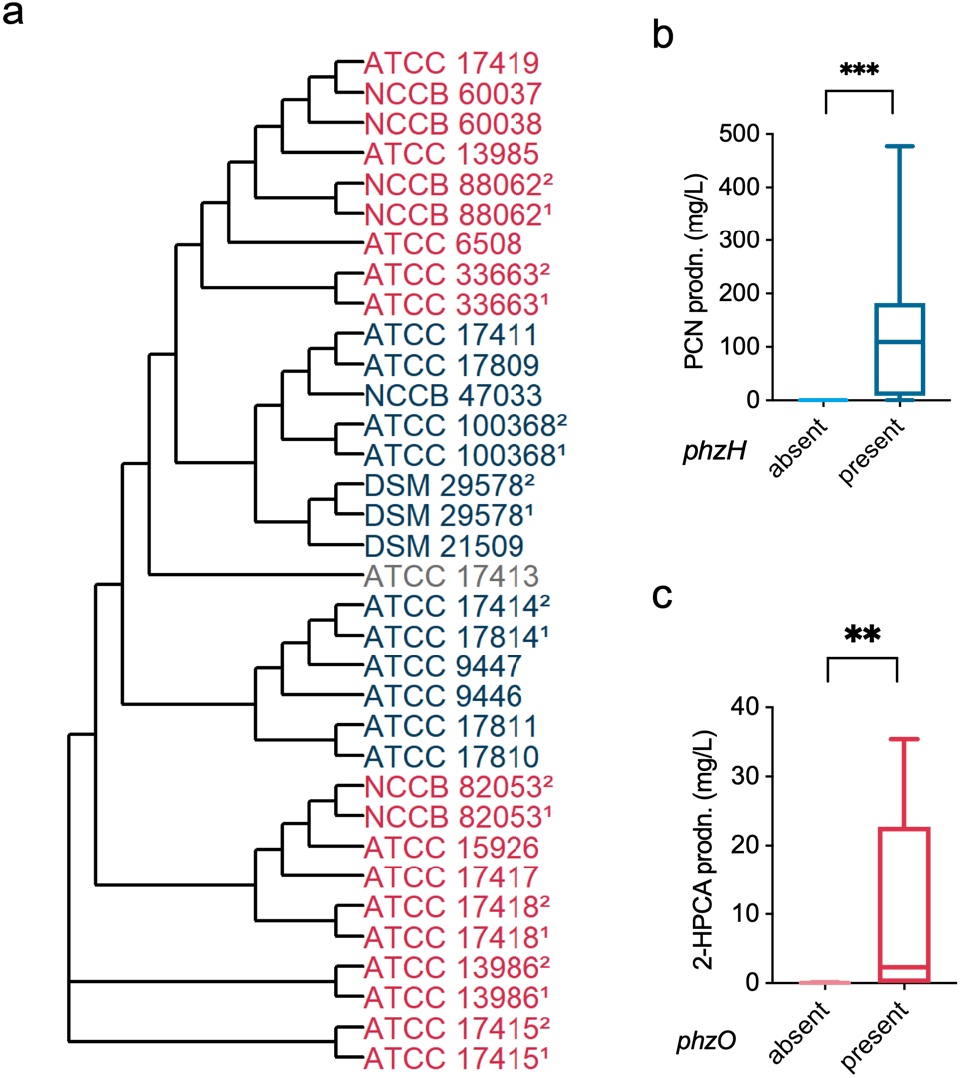
Categorizing isolates based on presence and absence of genes *phzH* and *phzO*. (a) Tree based on accessory gene content among the *P. chlororaphis* pangenome. Strains containing *phzO* (red) and *phzH* (blue) form groups with similar gene content. (b), (c) Box-and-whisker plots showing significantly higher (***p < 0.001) PCN production in strains containing *phzH* and significantly higher (**p < 0.01) 2-HPCA production in strains containing *phzO* in KMB+Fe based on paired t-test. Phenazine production data in (b) and (c) is taken from the phenotype data shown in Fig. 2.

### Identifying phenazine biosynthesis hits by mGWAS

We used the hybrid genome assemblies and the phenazine production data to carry out an mGWAS for identification of genetic signatures associated with phenazine production. Phenazine biosynthesis was split into a series of phenotypes pertaining to production of PCA, PCN, and total phenazines in KMB and KMB+Fe for the mGWAS analysis (see Materials and Methods for the complete list of phenotypes). **Table S3** shows the number of significant hits (*i*.*e*., unique DNA sequences or unitigs) obtained for each phenotype in the mGWAS. All hits and their reverse-complement sequences were aligned to genomes of all strains to find their genomic locations, resulting in a ‘preliminary list’ of 2493 significant hits across all phenotypes (**Fig. 5a, Supplementary File 1**). Each unitig (and its reverse-complement) in this list may be found in one or more strains and in one or more phenotypes, thus creating redundancies in the list as the unitigs were counted multiple times. These redundancies were eliminated by collapsing the hit list in 3 stages – phenotype-collapsing, strain-collapsing, and reverse-complement collapsing – to result in a ‘final list’ that only contains a unique entry for each significant hit influencing phenazine biosynthesis. Phenotype collapsing reduced the list to 1568; strain collapsing reduced the list further to 474. Finally, removing entries that were due to reverse complement redundancy resulted in a final list of 330 unique genomic hits for phenazine biosynthesis. The corrected p-value of the hits in the final list is shown in **Fig. 5b. Fig. S2** illustrates the collapsing pipeline, and **Supplementary File 2** contains all entries in each of the collapsed lists. A vast majority of the hits (284 out of 330) have a positive effect on phenazine production while the remaining ones had a negative effect (**Fig. S3**).

**Fig. 5.**
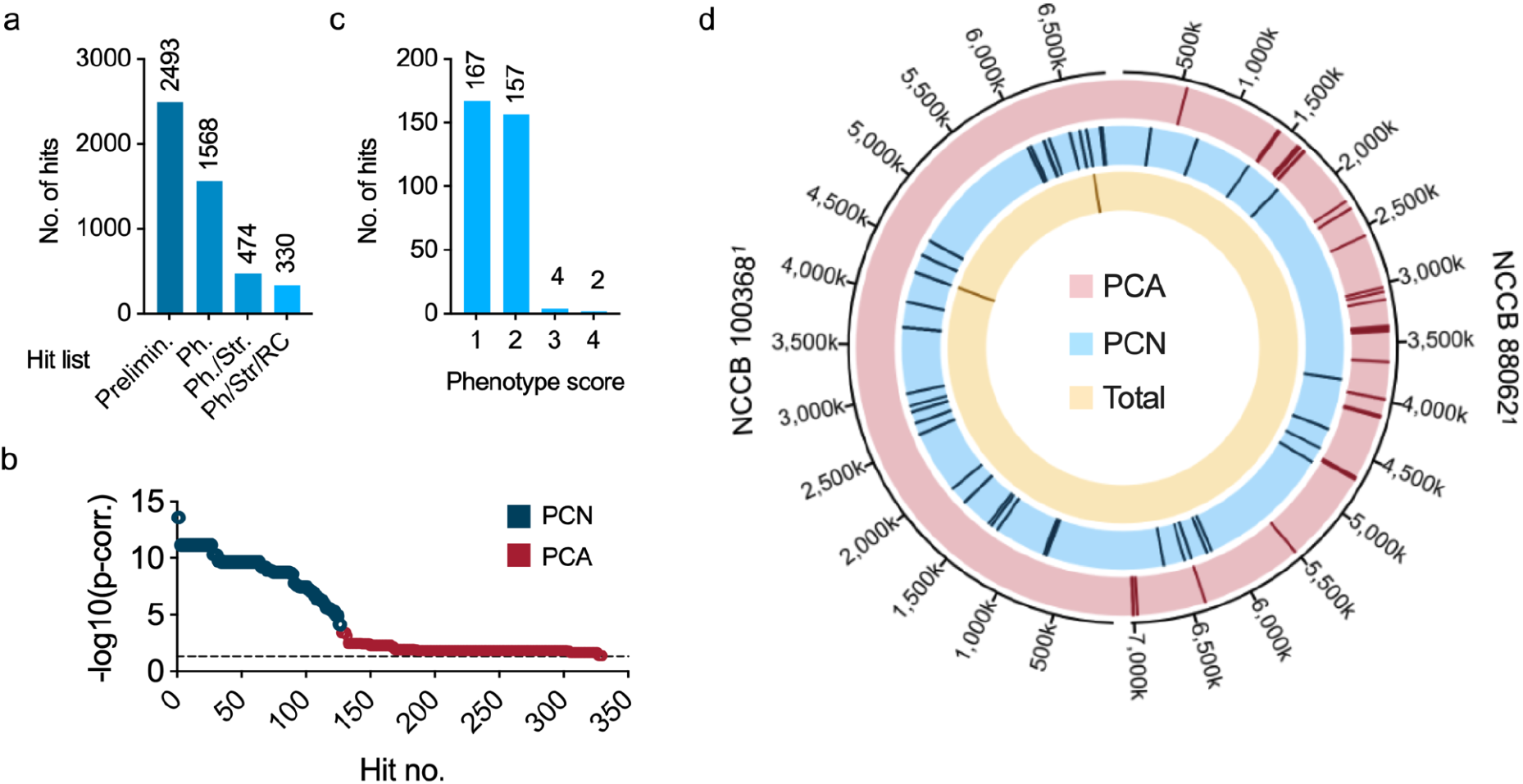
Results of the mGWAS analysis for phenazine production. (a) Number of significant mGWAS hits in the preliminary (uncollapsed) list and lists obtained after each collapsing stage - phenotype-collapsed list (Ph), phenotype+strain-collapsed list (Ph./Str.), and phenotype+strain+reverse complement-collapsed list (Ph/Str/RC; also called the ‘final list’). Numbers above each bar indicate the exact number of hits in the list corresponding to that bar. (b) Corrected p-values of the 330 hits in the final list. Hits were numbered in decreasing order of -log10(p-corrected) value, and were grouped into those influencing PCA production and PCN production. (c) Phenotype score distribution of hits in the final list. Numbers above each bar indicate the total number of hits having phenotype score corresponding to that bar. (d) Circos plot showing genomic locations of hits in the final list grouped into 3 categories based on the phenotype(s) in which they were significant: PCA production, PCN production, and total phenazine production, with respect to 2 strains - NCCB 100368^1^ and NCCB 88062^1^.

To visualize the genomic location of hits in the final list, we created a circos plot and mapped as many hits as possible to a PCN- or PCA-producing strain (**Fig. 5d**). Since each strain contains a different set of accessory genes and many of the hits map to the accessory genome, we were unable to map all hits to a single strain. We selected NCCB 100368^1^ as the basis for displaying the PCN-related hits because it contains the majority of the PCN hits (122 out of 127). Similarly, NCCB 88062^1^ was selected as the basis to display PCA related hits as it contained 170 out of 203 PCA-related hits, more than any other strain. In total, the two strains combined contain 292 hits from the final list (out of 330). Mapping the hits revealed that there is little association between the PCA hits and PCN-producing strains. In addition, the two hits related to total phenazine production mapped to PCN-producing strains only. In comparison, many of the hits related to PCN biosynthesis were found in both PCA- and PCN-producing strains.

The mGWAS analysis links unitigs to phenazine production, the next step in our analysis is to identify which genes are associated with each unitig. This mapping is straightforward when the unitig partially or completely overlaps to a CDS, but the connection to a gene is less clear when the unitig is contained within an intergenic region. In these cases, we identified genes upstream and downstream of the intergenic region as potential genes of interest. In total, the 330 unitig hits map to 158 genes in the pangenome (many of the genes were associated with more than one unitig; see **Supplementary File 3**). The 158 genes include 80 functionally annotated proteins or homologs of proteins with known function and 78 hypothetical proteins. While the function of the hypothetical proteins are unknown, 33 of them belonged to the core genome while the remainder belonged to the accessory genome. Forty-one out of the 80 proteins of known function belonged to the *P. chlororaphis* core genome.

In the phenotype-collapsing stage we associated each hit to a ‘phenotype score’, which represents the number of phenotypes in which a hit was found (**Fig. S2**); the higher the score, the greater the number of phenotypes influenced by the hit. The phenotype score was used as a metric to identify hits and associated genes most likely to improve phenazine biosynthesis. **Fig. 5c** shows the distribution of phenotype scores for the final list. We deemed hits with a phenotype score of 3 or higher as most likely to affect phenazine biosynthesis, and therefore ones that we sought to target for further analysis. The top phenazine-producing strain DSM 21509 primarily produces PCN, and the hits influencing PCN production have lower q-values than those influencing PCA production (Fig. 5b). We therefore narrowed our mGWAS validation and metabolic engineering studies to PCN producing hits only. **Table 2** shows the attributes of these hits, two of which have a phenotype score of 4, while the other two have a score of 3. All of these hits were found to be single nucleotide polymorphisms (SNPs) in the coding or intergenic (non-coding) regions. Genes containing or adjacent to SNPs for PCN production include YbhH (a putative isomerase), RhtA (threonine/homoserine exporter), UctC (acetyl-CoA:oxalate CoA-transferase), ProY_1 (proline-specific permease), HutH2 (histidine ammonia-lyase), and two hypothetical proteins (annotated as PS__04251 and PS__04252 in DSM 21509), all of which belong to the *P. chlororaphis* core genome. DSM 21509, the highest PCN-producing strain, contains 3 of the 4 SNPs for PCN production.

**Table 2.** Most influential hits for PCN production, identified in the final list of significant mGWAS hits. Hits are numbered 1 through 4, and attributes such as phenotype score, associated phenotypes, strains, and genes, as well as variant type, p-value, effect on phenotype (positive/negative), & genomic region have been provided for each hit.

| No. | Phenotype score | Major phenotypes | Associated gene(s) | Corr. p-value (effect) | Genomic region | Type of variant | Strains |
| --- | --- | --- | --- | --- | --- | --- | --- |
| 1 | 4 | PCN production<br>Total phz. production | hypothetical protein (PS_04251)<br>hypothetical protein (PS_04252) | 2.6 x 10<sup>-14</sup> (+) | intergenic | SNP | DSM 21509,<br>DSM 29578<sup>1</sup>,<br>DSM 29578<sup>2</sup>,<br>NCCB 100368<sup>1</sup> |
| 2 | 4 | PCN production<br>Total phz. production | Putative isomerase (YbhH) | 2.6 x 10<sup>-14</sup> (+) | CDS | SNP | DSM 21509,<br>DSM 29578<sup>1</sup>,<br>DSM 29578<sup>2</sup>,<br>NCCB 100368<sup>1</sup> |
| 3 | 3 | PCN production | Threonine/homoserine exporter (RhtA)<br>Acetyl-CoA:oxalate CoA-transferase (UctC) | 6.7 x 10<sup>-10</sup> (-) | intergenic | SNP | DSM 29578<sup>1</sup>,<br>DSM 29578<sup>2</sup>,<br>NCCB 100368<sup>2</sup> |
| 4 | 3 | PCN production | Proline-specific permease (ProY_1)<br>Histidine ammonia-lyase (HutH_2) | 2.2 x 10<sup>-9</sup> (+) | intergenic | SNP | DSM 21509,<br>DSM 29578<sup>1</sup>,<br>DSM 29578<sup>2</sup>,<br>NCCB 100368<sup>1</sup>,<br>NCCB 47033,<br>ATCC 17411,<br>ATCC 17809,<br>ATCC 17414,<br>ATCC 9446,<br>ATCC 17814,<br>ATCC 9447,<br>ATCC 17811,<br>ATCC 17810 |

### Validating mGWAS hits for PCN production

We overexpressed the top gene hits for PCN production in DSM 21509 to verify their phenotypic effects. Three of the genes yielded significant changes in PCN production when overexpressed in DSM 21509. Overexpression of PS__04252 in KMB produced 19.1 ± 3.5 mg/L of PCN compared to 58.7 ± 12.3 mg/L produced by the empty vector control, which is a 67.5% decrease (**Fig. 6a**). Overexpressing ProY_1 and PS__04251 increased PCN production in KMB+Fe to 420.2 ± 19.7 mg/L and 400.1 ± 21.5 mg/L, respectively, compared to the 343.6 ± 7.3 mg/L PCN produced by the empty vector control (**Fig. 6b**).

**Fig. 6.**
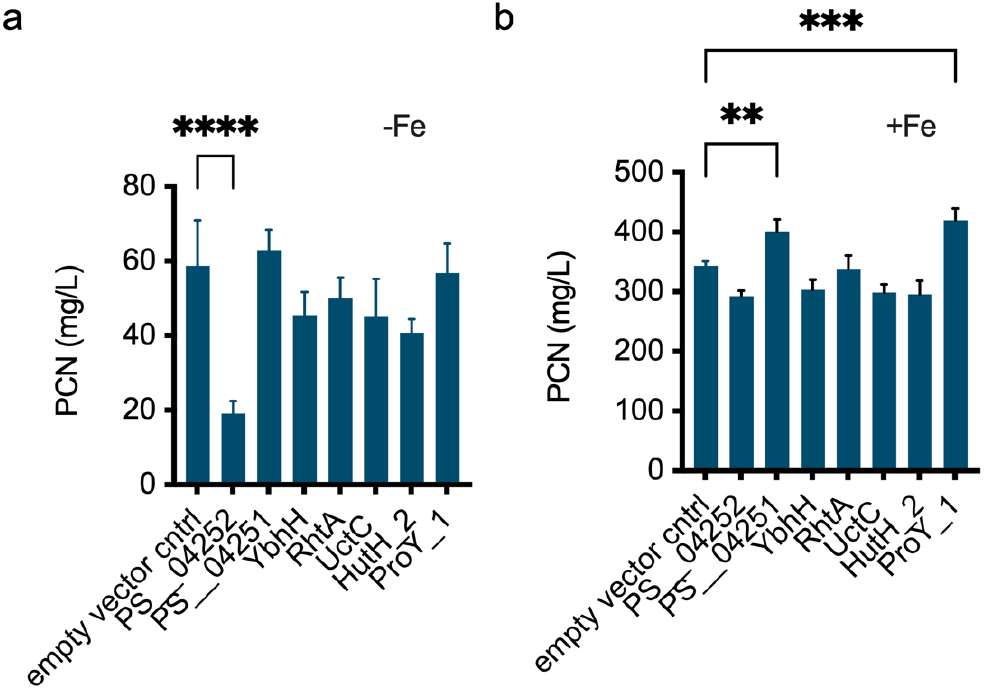
Overexpressing top hits in the top PCN-producer DSM 21509. Genes associated with top phenotype-scoring unitigs for PCN production were overexpressed in the top PCN-producing strain DSM 21509. PCN was quantified after 48 hr of culture in (a) King’s Media B and (b) King’s Media B + Fe. Bars represent the average of 3 replicates and error bars represent one standard deviation. Asterisks denote p-values <0.01 (**), <0.001 (***) and <0.0001 (****) when performing an ordinary one-way ANOVA comparison to the empty vector control.

PS__04251 and PS__04252 were both associated with unitig 1. PS__04252 is annotated as a hypothetical protein and has 57% sequence identity and 97% coverage to helix-turn-helix (HTH) MarR-family transcriptional regulator PA1607 from *P. aeruginosa* (NCBI Reference Sequence NP_250298.1) (Kaur and Subramanian, 2015). The MarR, or multiple antibiotic resistance repressor, family often regulates expression of multidrug efflux pumps and some, including PA1607, may derepress in response to oxidative stress (Housseini B Issa et al., 2018; Kaur and Subramanian, 2015). Therefore overexpression of the putative regulator PS__04252 could lead to increased repression of its target which could explain the observed decrease, rather than increase, in PCN production. PS__04251 is a hypothetical protein with unknown function which has 86% identity and 100% coverage to putative M14-type zinc cytosolic carboxypeptidase PSF113_3889 (NCBI Reference Sequence WP_041476041.1) from *Pseudomonas ogarae* whose function is unknown (Rimsa et al., 2014). While the function of this and similar proteins are unknown, we found its overexpression to significantly improve phenazine production in *P. chlororaphis*. Because both CDS surrounding unitig 1 were successful, this genomic region could be of further interest to investigate for phenazine production.

The other hit which significantly improved PCN production, ProY_1, was associated with unitig 4 which occurred within the hut operon, which is responsible for histidine catabolism. Due to its position in the highly conserved operon and its sequence similarity, the hit annotated as ProY_1 is likely the histidine permease HutT which imports histidine and is required for its utilization (Zhang et al., 2012). One study suggests that histidine catabolism in *P. fluorescens* is connected to oxidative stress response, as it could increase intracellular pools of the antioxidant α-ketoglutarate (Lemire et al., 2010). Possibly, overexpressing HutT could contribute to improved oxidative stress tolerance through a similar mechanism in *P. chlororaphis* by increasing intracellular histidine levels and therefore α-ketoglutarate levels. The other CDS adjacent to unitig 4 is HutH_2 (HutH2), the histidine ammonia lyase which catalyzes the first step of histidine catabolism, the conversion of L-histidine into urocanate (Zhang and Rainey, 2007). While this hit is associated with the same operon, its overexpression did not affect PCN titers. Because unitig 4 was within the non-coding region between 2 CDS, the mGWAS results in actuality may have been connected to one CDS and not the other. By overexpressing both hits, we were able to identify the one which was relevant for phenazine production.

While overexpressing the genes associated with unitigs 2 and 3 did not produce an observable change in phenotype, they also appear to be related to oxidative stress as well as amino acid export and catabolism. Unitig 2 occurred within the gene annotated as putative isomerase YbhH. This CDS has 34% identity and 92% coverage to *E. coli* YbhH, which does not have a known function but its expression has been upregulated in response to the *σ*^E^ stress signaling pathway (Bury-Moné et al., 2009). Unitig 3 occurred within a non-coding region which was flanked by 16s rRNA and either UctC or RhtA. The hit annotated as UctC has 87% sequence identity and 100% coverage to *P. putida* KT2440 glutarate-CoA transferase GcoT (NCBI Reference Sequence NP_742328.1) which is part of L-lysine catabolism (Zhang et al., 2019). The other associated hit has 57% identity and 93% coverage to *E. coli* RhtA (NCBI Reference Sequence WP_001295297), which exports threonine and homoserine in addition to other amino acids. While the mGWAS analysis identified these unitigs as significantly related to phenazine production, the overexpression studies showed that they did not alter PCN titers in DSM 21509.

In this study, we set out to identify novel genomic loci significant for the production of phenazine compounds in *P. chlororaphis*. We sequenced a collection of 34 *Pseudomonas* isolates, assembled their genomes and pangenome, and characterized 3 of their phenotypes (*i*.*e*., phenazine production, biofilm formation, and growth temperature) to select the best host strain. Through mGWAS analysis we identified 330 hits which were significant for phenazine production, and those hits mapped to 158 genes in the pangenome. We selected 7 genes associated with 4 of these hits and overexpressed them in DSM 21509, the strain with the highest PCN titers. We used the phenotype score to prioritize these hits over others because this method was unbiased in that it is blinded to gene function. Overall this data-driven approach was successful because we identified two candidate genes that improved phenazine production and one that reduced it.

An alternative approach is to prioritize hits based on a rational design strategy, that is, target genes with known functions related to the phenazine biosynthesis or associated pathways. For example, our hit list also includes GacA, a global transcriptional regulator known to impact phenazine production (Li et al., 2020; Wang et al., 2013). Pursuing this target or others associated with the Gac regulatory cascade could be promising for phenazine production, as the mGWAS and literature results are in agreement. Moreover, since we verified successful hits that may be transcriptional regulators or are involved in oxidative stress response, hits from the list with similar functions could be prioritized for future studies. The other hits on the list which are annotated as HTH-transcriptional regulators (*e*.*g*., ArgP, BenM, CynR, MtrA, and RhaS) or the hits known to be connected to oxidative stress response from the literature (*e*.*g*., RscC (Bury-Moné et al., 2009), glutathione synthase (Nikel et al., 2021; Wongsaroj et al., 2018) and glucose-6-phosphate 1-dehydrogenase (zwf) (Kim et al., 2008)) could be pursued as targets in future studies.

Many of the strains in our collection were isolated from diverse environmental locations before being deposited into their respective culture collections. Compared to a more traditional metabolic engineering approach that uses a single strain as a starting point to engineering, our population genomics approach takes advantage of natural phenotypic and genomic variation. In our analysis, we observed a broad range of phenotypes and genotypes that were distributed so that both positive and negative groups were well-represented. For example, similar numbers of strains contained *phzH* (14 strains) vs. *phzO* (19 strains), and similar numbers of strains produced no phenazines (10 strains), less than 100 mg/L phenazines (11 strains), and greater than 100 mg/L phenazines (13 strains). This phenotypic diversity, along with the genetic diversity found in the pangenome, allowed us to use a relatively small number of strains to perform the mGWAS analysis and obtain significant hits. The statistical power of hit identification can be potentially improved by using a bigger and more diverse collection of isolates as input to the mGWAS analysis, entailing a larger representation of phenotypic and genomic diversity. Further, GO and pathway enrichment tests of genes associated to 330 hits resulted in no enriched terms (see **Supplementary File 4, Supplementary File 5**). Pursuing hits based off their functions will become more promising as more microbes are annotated for GO-terms and metabolic pathways.

Taken together, this work presents a new approach that enables genome-scale metabolic engineering of pseudomonads. This approach is data-driven, using the power of low cost sequencing and high throughput phenotyping to generate large data sets that correlate desired traits to genomic variants within a microbial population, thereby generating new metabolic engineering targets. Here we show one example of this approach; we anticipate that it could be used in future studies to further enhance biosynthesis of other natural products or to improve other industrially relevant phenotypes in pseudomonads or in other microbial hosts.

## Conclusions

Advancements in whole genome DNA sequencing and genome-editing techniques, as well as increased availability of bioinformatics tools for analysis of genome-wide data have allowed us to identify metabolic engineering targets spanning the entire pangenome. The accessory genome and core genome are promising sources of metabolic engineering targets for the bacterial production of secondary metabolites such as phenazines. The present study taps into both of these pangenome components to help identify strain engineering targets for biosynthesis of the phenazine PCN in *Pseudomonas chlororaphis*. This pangenome-wide approach, in combination with rational design approaches, could potentially lead to substantial improvement in the phenotype of interest, while also assisting with selection of the appropriate host strain for metabolic engineering.

## Materials and methods

### Strain selection and culturing

All strains designated as *Pseudomonas chlororaphis* that were available as of April and October 2019 were ordered from the American Type Culture Collection (ATCC; Manassas, VA); all strains designated as *Pseudomonas chlororaphis* that were nonredundant and available as of March 2020 were ordered from the German Collection of Microorganisms and Cell Cultures (DSMZ GmbH; Braunschweig, Germany) and the Westerdijk Fungal Biodiversity Institute’s Netherlands Culture Collection of Bacteria (NCCB; Utrecht, Netherlands). Strains which appeared to have more than one colony morphology present were separated into distinct isolates (denoted by ^1^ and ^2^, which were arbitrarily assigned) that were sequenced and cultured separately. All isolates were sequenced with 16s rRNA sequencing (GENEWIZ®; South Plainfield, NJ), and the 33 confirmed *P. chlororaphis* isolates were used in this study (**Table S1**). One PCA-producing *P. synxantha* isolate was also used in this study as a phylogenetic outgroup for a total of 34 isolates.

Strains were initially revived according to the guidance of each culture collection then subsequently cultured at 30°C in King’s Media B (KMB), the standard media for fluorescent pseudomonads culture, according to the methods of King et al. (King et al., 1954). To improve phenazine production, KMB+Fe Media was made by supplementing KMB with 100 μM ferric sodium ethylenediaminetetraacetate (FeNaEDTA) based off the findings of van Rij et al. (van Rij et al., 2004). For phenazine production experiments, both KMB and KMB+Fe contained 1.5 g/L MgSO_4_. Cultures were supplemented with 50 μg/mL kanamycin sulfate when an antibiotic resistance marker was used. Luria Bertani (LB) broth and TOP10 chemically competent *E. coli* cells were used for cloning.

Liquid culturing was performed using sterile 2 mL 96-deep well plates within an INFORS HT Multitron Pro plate shaker incubator at 1000 rpm and ∼88% humidity. Overnight cultures were started by inoculating 500 μL media of interest with the respective colony or glycerol stock. After the overnight culture incubated with shaking at 30 °C for 22-24 hours, the plate was spun down in Beckman Coulter Allegra 25R centrifuge for 10 minutes at 5,000g. To reduce phenazine transfer and to ensure biofilm-forming strains were well-mixed, old media was removed, and cultures were resuspended in fresh media. To start experimental cultures, 500 μL of desired media were inoculated with 10 μL of resuspended overnight culture. For cultures requiring induction with isopropyl β-D-1-thiogalactopyranoside (IPTG), sterile-filtered IPTG was added to cultures to a final concentration of 1 mM about 4 hours after inoculation.

### Phenazine quantification

48 hours after inoculation, phenazine compounds were extracted from each liquid culture using ethyl acetate liquid-liquid extraction. Whole cultures were acidified with 10 μL of 3M HCl, then 1.2 mL ethyl acetate was added to each culture. Each mixture was transferred to a microcentrifuge tube, vortexed at maximum speed for 1 minute, and spun down to separate liquid phases. The ethyl acetate phase was evaporated, resuspended in methanol, and filtered for quantification via HPLC.

All phenazines were quantified with a photodiode-array detector on a Shimadzu Nexera-i LC-2040C 3D liquid chromatograph with an Agilent Poroshell 120 EC-C18 2.7 μm 3.0 × 75 mm column and 3.0 mm x 5.0 mm guard column at 40°C. To resolve the similar phenazine derivatives, the following method with gradients of methanol and ammonium acetate buffer (pH 5.0) was used: 2 μL sample injection, 5 min of 20% methanol, 2 min gradient from 20% to 30% methanol, and 13 min gradient from 30 to 40% methanol with subsequent steps to wash and re-equilibrate the column, with all steps at a 1 mL/min flow rate. PCA and PCN peaks were identified by comparing retention times to those of purchased PCA and PCN (ChemScene; Monmouth Junction, NJ). Because pure 2-HP and 2-HPCA were not commercially available, the identities of these HPLC peaks were confirmed with LC-MS following the same protocol. Phenazines were quantified by converting peak areas at a wavelength of 254 nm and bandwidth of 4 nm to concentrations using extinction constants calculated from the purchased PCA and PCN.

### Biofilm formation and growth temperature phenotyping

Biofilm formation phenotyping was characterized using a crystal violet staining assay following the protocol of O’Toole (O’Toole, 2011). To adapt the protocol for *Pseudomonas chlororaphis*, a 2% inoculum of overnight culture in the respective media was used to start stationary cultures which were incubated without agitation at 30 °C for 48 hours. Growth temperature phenotyping was performed by streaking overnight culture in patches on the respective solid media and incubating the plates at either 30 °C or 37 °C for 48 hours. Plates were assessed visually for growth based on the density of the culture patch.

### Genome assembly

Genomic DNA was isolated using the Quick-DNA™ Fungal/Bacterial Miniprep Kit (Zymo Research; Irvine, CA) and sent to the Microbial Genome Sequencing Center (Pittsburgh, PA) for whole genome sequencing. All isolates were sequenced on the NextSeq 2000 (Illumina; San Diego, CA) with paired-end 150 base pair reads and with Oxford Nanopore technologies. Illumina read quality was assessed using FASTQC v0.11.9 (Andrews and Others, 2010) and Nanopore read statistics were assessed with NanoStats v1.28.2 (De Coster et al., 2018) on the Galaxy platform before and after read filtering and trimming (the parameters for all bioinformatics tools are available in **Table S2**). Summary statistics (*i*.*e*., total assembly length, number of contigs, N_50_, L_50_, % GC) were calculated using QUAST v.5.0.2 (Mikheenko et al., 2018). Genome completeness was assessed by running BUSCO v.5.2.2 in genome mode using the pseudomonadales_odb10 (prokaryota, 2020-03-06) database (Manni et al., 2021).

Flye genome assemblies were created with Flye v.2.8.3 and raw Nanopore reads as input (Kolmogorov et al., 2019). All other genome assemblies used reads which were filtered and trimmed based on read quality. Raw Illumina reads were trimmed to remove adapters and low-quality ends using Trimmomatic v.0.38 (Bolger et al., 2014). Raw Nanopore reads were adaptor-trimmed using Porechop v.0.2.4 (Wick, 2018) then filtered with filtlong v.0.2.1 (Wick and Menzel, 2019). SPAdes genome assemblies were created using SPAdes v.3.12.0 (Prjibelski et al., 2020) on the Galaxy platform (Afgan et al., 2018); Unicycler assemblies were created using Unicycler v.0.4.8 using both “Normal” and “Bold” bridging modes and excluding contigs shorter than 1000 bp from the assemblies (Wick et al., 2017). The short-reads assemblies were created using only the paired end Trimmomatic output. The long-reads Nanopore assemblies were created using the trimmed and filtered Nanopore reads. The hybrid assemblies were assembled using both sets of aforementioned reads.

### Genome annotation and pangenome assembly

Genome assemblies were annotated with Prokka v1.14.6 (Seemann, 2014) on the Galaxy platform (Afgan et al., 2018), using a minimum contig size of 1000 and ‘Pseudomonas’ as the genus name. Prokka outputs genome annotations in GFF3 format. These GFF3 files were used along with the draft genome assemblies to generate annotated genome FASTA files by bedtools GetFastaBed v2.30.0 (Quinlan and Hall, 2010). The pangenome was constructed using PEPPAN v1.0.5 and the .gff files generated by Prokka as input (Zhou et al., 2020). A rarefaction curve, gene presence absence matrix, and accessory genome tree were created from the PEPPAN output using the included PEPPAN_parser algorithm. Statistics about the core genome were calculated from the gene presence absence matrix using R v4.2.1 (RStudio 2022.07.1). The resulting .nwk tree file was visualized using R v4.2.1 and treeio package v1.20.2 (Wang et al., 2020). All remaining figures were created using GraphPad Prism v9.4.1 (GraphPad Software; San Diego, CA).

### Genome-wide association study

De novo assembled genomes of the 34 *Pseudomonas* strains were provided as input to DBGWAS v0.5.4 (Jaillard et al., 2018) along with the corresponding phenotype values for phenazine production. DBGWAS was implemented for 7 different phenotypes: (i) PCA production in KMB; (ii) PCA production in KMB+Fe; (iii) Effect of Fe on PCA production; (iv) PCN production in KMB; (v) PCN production in KMB+Fe; (vi) Effect of Fe on PCN production; (vii) Total phenazine production in KMB.

For phenotypes (i), (ii), (iv) and (v), concentrations of PCA and PCN (mg/L) obtained from experiments were used directly. Values of phenotypes (iii) and (vi) were estimated by subtracting the concentration of the phenazine compound in KMB from that in KMB+Fe. If the difference was negative, it was replaced by 0. Total phenazine production in KMB was obtained by simply adding the concentrations of all phenazine compounds (*i*.*e*., PCA, 2-HPCA, PCN and 2-HP) in KMB. The genome sequences of strains were also provided as BLAST database to DBGWAS for genome mapping of significant unitigs. Significant unitigs were identified based on a corrected p-value cutoff, and a minor allele frequency greater than 1% (default). The number of significant unitigs obtained for each phenotype are listed in **Table S3**.

### Downstream processing of mGWAS hits

Even though DBGWAS maps significant unitigs to genomes by BLAST, we chose to independently perform unitig alignment to genomes by exact matching to avoid any tolerance to mismatches during alignment. Custom Python3 scripts were used for this purpose with the 34 genome sequences as the mapping database. To ensure that each unitig finds a match, both the significant unitig and its reverse-complement were used. Further, genome annotations were used to determine the genomic regions of the mapped unitigs (*i*.*e*., whether the unitig falls within a gene or an intergenic region).

The lists of mGWAS hits from the 7 phenotypes were concatenated into a single list called the ‘preliminary list’. In this list, each occurrence of a significant unitig constituted a single entry, creating separate entries for each phenotype, strain, as well as the reverse complement sequence of that unitig. Custom Python3 scripts were used to remove redundancies and collapse the preliminary list so that each significant unitig has a single entry in the final list (**Fig. S2**) For each strain, entries for identical unitigs that were significant for multiple phenotypes were collapsed together in the ‘phenotype-collapsed’ list. Each entry was assigned a phenotype score, which represents the number of phenotypes (out of 7) where each unitig was significant. If a unitig had a phenotype score greater than 1, its corrected p-value was taken to be the minimum of corrected p-values for all phenotypes it is found in. Similarly, the effect of that unitig was taken to be the one with the highest magnitude across all phenotypes. The ‘phenotype+strain-collapsed list’ combined entries where the same unitig mapped to the same genomic region in multiple strains. Redundancies where the reverse complement of a significant unitig shows up as a separate entry were then collapsed to create the ‘final list’ of mGWAS hits.

Genes associated to mGWAS hits were determined based on the overlap of unitigs to genes. If the overlap to a gene was partial or complete, that gene was considered to be associated to the unitig. In case of no overlap, i.e., when the unitig appeared completely between two genes, both the neighboring genes were considered to be linked to the unitig.

### Experimental validation of top hits

The hits from the final list with a phenotype score of 3 or higher which were significant for PCN production phenotypes were selected as top hits for experimental validation. The CDS immediately upstream and downstream of each significant unitig were chosen as metabolic engineering targets to be overexpressed in the top PCN-producing strain. Any CDS which encoded ribosomal RNA was discarded from the list. If the significant unitig sequence was completely contained within a CDS, only the unitig-containing CDS was studied rather than the 2 adjacent CDS.

Each target was PCR-amplified from the genomic DNA of the strain listed on the top hits file then inserted into the backbone of plasmid pBb(RK2)1k-GFPuv using either restriction digest cloning or NEBuilder® HiFi DNA Assembly (New England Biolabs; Ipswich, MA) using the primers listed in **Table S4**. pBb(RK2)1k-GFPuv is a broad-range expression vector with an IPTG-inducible promoter which was gifted by Brian Pfleger at the University of Wisconsin, Madison, and used as the empty vector control (Cook et al., 2018). All plasmids used in this study are listed in **Table S5**.

Plasmids were transformed into the respective strain via electroporation based on the method of Choi et al. (Choi et al., 2006). Electroporations were performed by pulsing either 1.8 or 2.5 kV through a 0.1 or 0.2 cm electroporation cuvette using a MicroPulser™ Electroporation Apparatus (Bio-Rad) then recovering the reaction for 2-3 hours at 30°C. Culturing and phenazine quantification were performed as described in previous subsections.

### Gene Ontology enrichment analysis

To identify enriched GO-terms for significant mGWAS hits, strain DSM 21509 (highest PCN producer) was used as the reference. GO-IDs for this strain were obtained using Blast2GO v6.0.3 (Conesa et al., 2005). First, Blast2GO was used to map the annotated genome of strain DSM 21509 to proteins in the *P. chlororaphis* protein file (program: blastx; number of blast hits = 5; HSP length cutoff = 50) obtained from NCBI (Taxonomy ID: 587753). Next, the BLAST hits were mapped to GO-identifiers from the database of the Gene Ontology Consortium (Ashburner et al., 2000; Harris et al., 2004). Lastly, GO mapped hits were annotated (Hit Filter = 2; Filter GO by taxonomy: g-proteobacteria (taxa: 1236,Gammaproteobacteria)) to obtain additional information, such as enzyme codes, enzyme names and InterPro IDs. GO-enrichment test was performed with the obtained GO-IDs using the tool GOEnrichment v2.0.1 on the Galaxy platform (Afgan et al., 2018). Annotated genes associated to significant unitigs in the final list were provided as the study set. GO annotations from the strain DSM 21509 were provided as the reference set. All enrichment tests were performed using an FDR-corrected p-value cutoff of 0.05 for enrichment.

### Pathway enrichment analysis

A list of existing metabolic pathways (and corresponding genes involved) in *P. chlororaphis* strain PA23 was extracted from KEGG PATHWAY database (Kanehisa et al., 2017) (prefix: pch) and written into a custom pathway database file (.GMT format) using R 4.2.1 (RStudio 2022.07.1). Sequences of all genes associated to significant unitigs in the final list were BLASTed against proteins of the *P. chlororaphis* strain PA23 (obtained from NCBI) using Blast2GO v6.0.3 (Conesa et al., 2005) (program: blastx; number of blast hits = 5; HSP length cutoff = 50) to find homologs. The list of PA23 gene homologs was then used as input along with the custom pathway database file to perform pathway enrichment analysis using the web version of the tool g:Profiler (Raudvere et al., 2019).

## Abbreviations

CDS: coding sequence(s)
KMB: King’s media B
KMB+Fe: King’s media B supplemented with ferric iron
mGWAS: microbial genome-wide association studies
PCA: phenazine-1-carboxylic acid
PCN: phenazine-1-carboxamide
2-HP: 2-hydroxyphenazine
2-HPCA: 2-hydroxyphenazine-1-carboxylic acid

## Data availability

Sequencing reads and assembled genomes for the 34 *Pseudomonas* isolates have been deposited in the NCBI SRA (BioProject ID: PRJNA932460) and NCBI GenBank databases, respectively. NCBI accession numbers for the assembled genomes have been provided in Table 1. Scripts used for collapsing mGWAS hits have been provided as supplementary material (**Supplementary File 6**).

## Author contributions

ST, VT and IW conceived the project, planned the experiments, and analyzed the data. ST collected the bacterial isolates, sequenced and assembled the whole genomes and the pangenome, and conducted the phenotype screening experiments for the isolates. VT implemented the mGWAS analysis and performed downstream processing of mGWAS hits. ST carried out validation experiments for the top mGWAS hits. All authors wrote and edited the manuscript.

## Acknowledgements

This work was supported by the Army Research Office MURI (#W911NF1410263), Air Force Office of Scientific Research award FA9550-17-1-0270, and NSF Plants-3D 1922642.

