## Supplementary Information for "Population genomics-guided engineering of phenazine biosynthesis in *Pseudomonas chlororaphis*"

Fig. S1: Graphical comparisons of the summary statistics for all genome assemblies.

Fig. S2: Collapsing pipeline for GWAS hits.

Fig. S3: Distribution of significant GWAS hit effects in the ‘final list’.

Table S1: List of strains used in this study.

Table S2: List of bioinformatics tools and parameters used for genome and pangenome assembly and annotation in this study.

Table S3: Number of significant unitigs obtained for different phenotypes in the GWAS analysis.

Table S4: List of primers used in this study.

Table S5: List of plasmids used in this study.

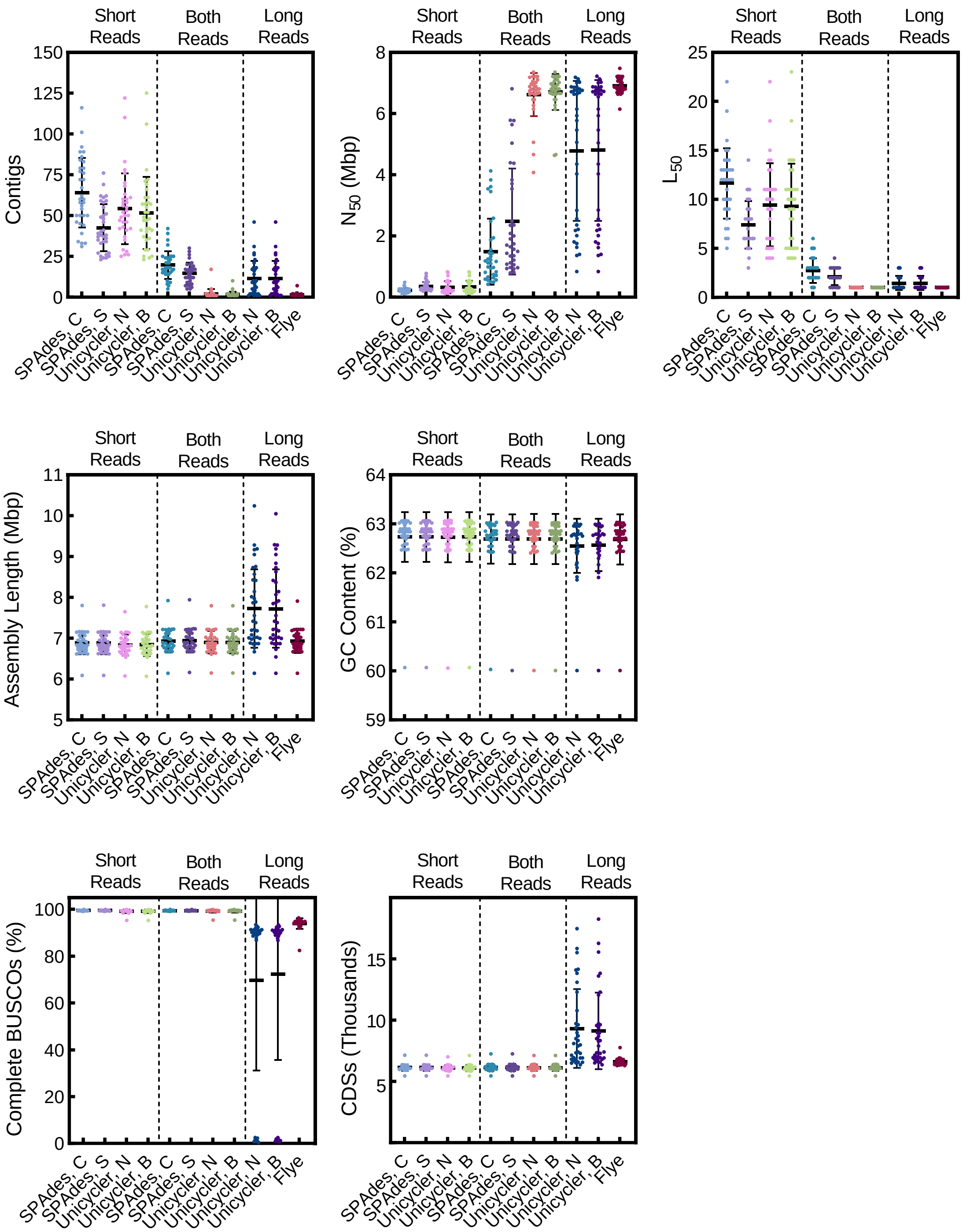


**Figure S1. Graphical comparisons of the summary statistics for all genome assemblies.** Genome assemblies were created with only short Illumina sequencing reads (left graph subsections), only long Oxford Nanopore sequencing reads (right graph subsection) or both reads sets in a hybrid approach (middle graph subsection), with contigs files (C) and scaffolds (S) generated by SPAdes, and genome assemblies generated by using either normal (N) or bold (B) bridging mode in Unicycler. Number of contigs, $N_{50}$, $L_{50}$, total sequence length, and GC content were generated using QUAST. The % complete BUSCOS was calculated using BUSCO and number of CDS from assembly annotations generated by Prokka

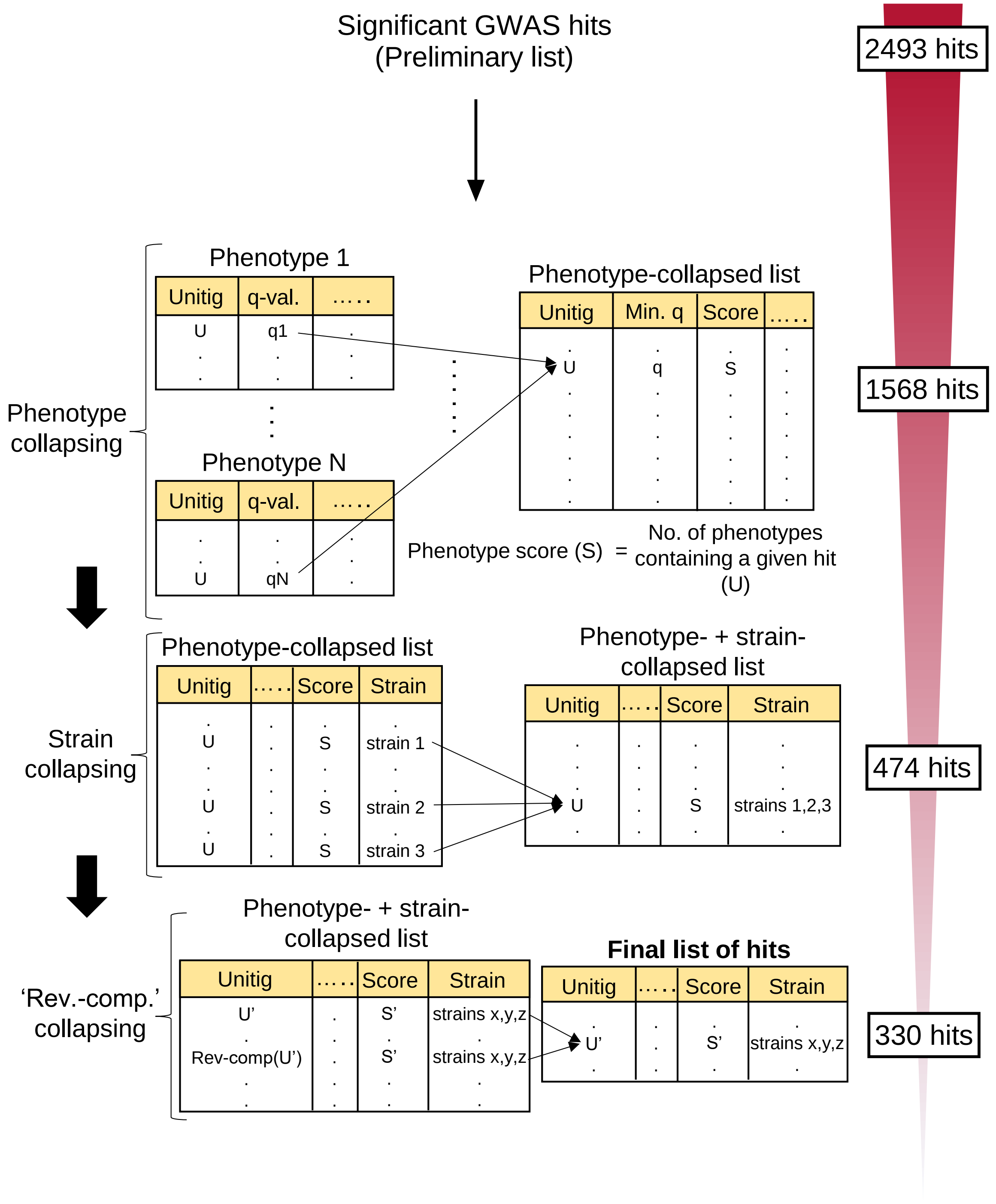


**Figure S2. Collapsing pipeline for GWAS hits.** A schematic showing each collapsing stage to remove redundancies in the preliminary list of significant GWAS hits. A phenotype score is assigned to each hit in the phenotype-collapsing stage. Numbers on the right indicate the total number of hits in the preliminary list as well as resulting lists from each corresponding collapsing stage for the phenazine production GWAS results.

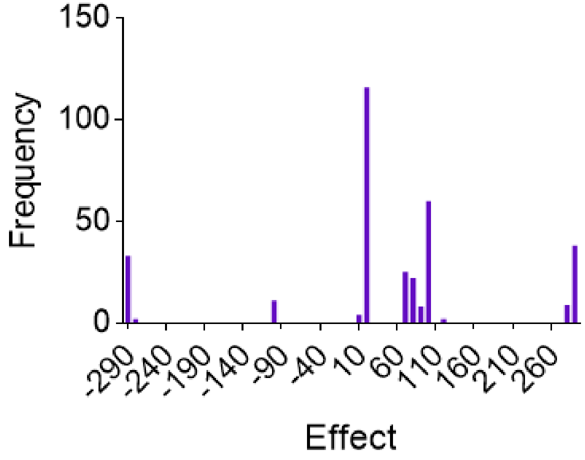


**Figure S3. Distribution of significant GWAS hit effects in the 'final list'.** Vertical axis indicates the number of significant hits having effect values in a certain range as specified by each bin.

**Table S1. List of strains used in this study.** All of the listed isolates were 16s rRNA sequenced and confirmed to be *Pseudomonas chlororaphis* except for ATCC 17413 which was determined to be *Pseudomonas synxantha*.

| **Full strain name** | **Strain isolate used in this study** | **Source** |
|---|---|---|
| *Pseudomonas chlororaphis subsp. chlororaphis* (ATCC® 9446™) | ATCC 9446 | American Type Culture Collection |
| *Pseudomonas chlororaphis* (ATCC® 9447™) | ATCC 9447 | American Type Culture Collection |
| *Pseudomonas chlororaphis subsp. aureofaciens* (ATCC® 13985™) | ATCC 13985 | American Type Culture Collection |
| *Pseudomonas chlororaphis* (ATCC® 13986™) | ATCC 13986[1]<br>ATCC 13986[2] | American Type Culture Collection |
| *Pseudomonas chlororaphis* (ATCC® 15926™) | ATCC 15926 | American Type Culture Collection |
| *Pseudomonas chlororaphis* (ATCC® 17411™) | ATCC 17411 | American Type Culture Collection |
| *Pseudomonas chlororaphis* (ATCC® 17413™) | ATCC 17413 | American Type Culture Collection |
| *Pseudomonas chlororaphis* (ATCC® 17414™) | ATCC 17414 | American Type Culture Collection |
| *Pseudomonas chlororaphis* (ATCC® 17415™) | ATCC 17415[1]<br>ATCC 17415[2] | American Type Culture Collection |
| *Pseudomonas chlororaphis* (ATCC® 17417™) | ATCC 17417 | American Type Culture Collection |
| *Pseudomonas chlororaphis* (ATCC® 17418™) | ATCC 17418[1]<br>ATCC 17418[2] | American Type Culture Collection |
| *Pseudomonas chlororaphis* (ATCC® 17419™) | ATCC 17419 | American Type Culture Collection |
| *Pseudomonas chlororaphis* (ATCC® 17809™) | ATCC 17809 | American Type Culture Collection |
| *Pseudomonas chlororaphis* (ATCC® 17810™) | ATCC 17810 | American Type Culture Collection |
| *Pseudomonas chlororaphis* (ATCC® 17811™) | ATCC 17811 | American Type Culture Collection |
| *Pseudomonas chlororaphis* (ATCC® 17814™) | ATCC 17814 | American Type Culture Collection |
| *Pseudomonas chlororaphis subsp. aurantiaca* (ATCC® 33663™) | ATCC 33663[1]<br>ATCC 33663[2] | American Type Culture Collection |
| *Pseudomonas chlororaphis subsp. aureofaciens* DSM-6508 | DSM 6508 | German Collection of Microorganisms and Cell Cultures |
| *Pseudomonas chlororaphis subsp. piscium* DSM-21509 | DSM 21509 | German Collection of Microorganisms and Cell Cultures |
| *Pseudomonas chlororaphis subsp. aureofaciens* DSM-29578 | DSM 29578[1]<br>DSM 29578[2] | German Collection of Microorganisms and Cell Cultures |
| *Pseudomonas chlororaphis* subsp. *chlororaphis* NCCB 47033 | NCCB 47033 | Netherlands Culture Collection of Bacteria |
| *Pseudomonas chlororaphis* subsp. *aureofaciens* NCCB 60037 | NCCB 60037 | Netherlands Culture Collection of Bacteria |
| *Pseudomonas chlororaphis* subsp. *aureofaciens* NCCB 60038 | NCCB 60038 | Netherlands Culture Collection of Bacteria |
| *Pseudomonas chlororaphis* subsp. *aureofaciens* NCCB 82053 | NCCB 82053[1]<br>NCCB 82053[2] | Netherlands Culture Collection of Bacteria |
| *Pseudomonas chlororaphis* subsp. *aureofaciens* NCCB 88062 | NCCB 88062[1]<br>NCCB 88062[2] | Netherlands Culture Collection of Bacteria |
| *Pseudomonas chlororaphis* subsp. *chlororaphis* NCCB 100368 | NCCB 100368[1]<br>NCCB 100368[2] | Netherlands Culture of Collection Bacteria |

**Table S2. List of bioinformatics tools and parameters used for genome and pangenome assembly and annotation in this study.** Other parameters which are not listed were kept at default values.

| Tool Name | Version | Parameters |
|---|---|---|
| FASTQC | v0.11.9 | Input: Illumina reads, raw and output from Trimmomatic (forward and reverse fastq.gz files)<br>• Default settings |
| NanoStats | v1.28.2 | Input: Oxford Nanopore reads, raw and output from Porechop and filtlong (fastq.gz files)<br>• Default settings |
| QUAST | v5.0.2 | Input: All genome assemblies (fasta files)<br>• Default settings |
| BUSCO | v5.2.2 | Input: All genome assemblies (fasta files)<br>• Lineage dataset (--lineage-dataset): pseudomonadales_odb10 (prokaryota, 2020-03-06)<br>• Running mode (--mode): genome |
| Flye | v2.8.3 | Input: raw Oxford Nanopore reads (fastq.gz files)<br>• Specify raw Oxford Nanopore reads as input (--nano-raw) |
| Trimmomatic | v0.38 | Input: raw Illumina reads (forward and reverse fastq.gz files)<br>• ILLUMINACLIP: NexteraPE<br>• LEADING:3<br>• TRAILING:3<br>• SLIDINGWINDOW: 4:15<br>• MINLEN:36 |
| Porechop | v0.2.4 | Input: raw Oxford Nanopore reads (fastq.gz files)<br>• Default settings |
| filtlong | v0.2.1 | Input: adapter-trimmed Oxford Nanopore reads output from Porechop (fastq.gz files)<br>• Minimum length (--min_length): 1000<br>• Minimum mean quality (--min_mean_q): 10 |
| SPAdes | v3.12.0 | Input: paired-end trimmed Illumina reads output from Trimmomatic (forward and reverse fastq.gz files). Hybrid assemblies also used filtered long reads output from filtlong (fastq.gz files)<br>• Default settings |
| Unicycler | v0.4.8 | Input: paired-end trimmed reads output from Trimmomatic. Hybrid assemblies also used filtered long reads output from filtlong.<br>• Bridging mode (--mode): normal OR bold<br>• Exclude contigs shorter than this length from FASTA (bp) (--min_fasta_length: 1000 |
| Prokka | v1.14.6 | Input: All genome assemblies (fasta files)<br>• Default settings |
| PEPPAN | v1.0.5 | Genome annotations output from Prokka (.gff files for either all final genome assemblies or only the *P. chlororaphis* assemblies) |
| PEPPAN_parser | | Output by PEPPAN output (PEPPAN.gff file)<br>• Generate gene presence/absence tree (--tree)<br>• Generate rarefaction curve (--curve)<br>• Ignore pseudogenes in analysis (--pseudogene). This flag was used in the pangenome analysis, but left of when determining which hits belonged to the core and accessory genome |
| treeio | v1.20.2 | Input: gene presence/absence tree output by PEPPAN_parser (All 34 strains, excluding pseudogenes; CDS_content.nwk file) |

**Table S3. Number of significant unitigs obtained for different phenotypes in the GWAS analysis.**

| Phenotype | No. of significant unitigs |
|---|---|
| PCA production in KMB | 121 |
| PCA production in KMB + Fe | 77 |
| Effect of Fe on PCA production | 79 |
| PCN production in KMB | 122 |
| PCN production in KMB + Fe | 81 |
| Effect of Fe on PCN production | 4 |
| Total phenazine production in KMB | 2 |

**Table S4. List of primers used in this study.**

| Name | Sequence (5’→3’) | Description |
|---|---|---|
| PS__04252_F | AATTTCAGAATTCAAAAGATCTTTTAAGAAGGAGATATACATATGGTCAAACGCACAAGC | Amplify PS__04252 from DSM 21509 gDNA for restriction digest cloning into pBb(RK2)1k-GFPuv backbone, forward primer |
| PS__04252_R | CCTTACTCGAGTTTGGATCCCTATTGCACCGGCACCC | Amplify PS__04252 from DSM 21509 gDNA for restriction digest cloning into pBb(RK2)1k-GFPuv backbone, reverse primer |
| PS__04251_F | AATTTCAGAATTCAAAAGATCTTTTAAGAAGGAGATATACATATGACCGTGGCTCAAAGC | Amplify PS__04251 from DSM 21509 gDNA for restriction digest cloning into pBb(RK2)1k-GFPuv backbone, forward primer |
| PS__04251_R | ATCCTTACTCGAGTTTGGATCCTCAGCGCAGGATGCCGA | Amplify PS__04251 from DSM 21509 gDNA for restriction digest cloning into pBb(RK2)1k-GFPuv backbone, reverse primer |
| RhtA_F | AATTTCAGAATTCAAAAGATCTTTTAAGAAGGAGATATACATATGAATGACCAGCCCCG | Amplify *rhtA* from DSM 29578[2] gDNA for restriction digest cloning into pBb(RK2)1k-GFPuv backbone, forward primer |
| RhtA_R | ATCCTTACTCGAGTTTGGATCCTCAATCAGCTGCAACCAAAG | Amplify *rhtA* from DSM 29578[2] gDNA for restriction digest cloning into pBb(RK2)1k-GFPuv backbone, reverse primer |
| ProY1_F | AATTTCAGAATTCAAAAGATCTTTTAAGAAGGAGATATACATATGCAACAGCAAGCTCAA | Amplify *proY_1* from ATCC 9447 gDNA for restriction digest cloning into pBb(RK2)1k-GFPuv backbone, forward primer |
| ProY1_R | ATCCTTACTCGAGTTTGGATCCTTATCGATGGGACAAAGAAGG | Amplify *proY_1* from ATCC 9447 gDNA for restriction digest cloning into pBb(RK2)1k-GFPuv backbone, reverse primer |
| UctC_F | AATTTCAGAATTCAAAAGATCTTTTAAGAAGGAGATATACATATGGGCGCGTTATCTCAT | Amplify *uctC* from DSM 29578[2] gDNA for restriction digest cloning into pBb(RK2)1k-GFPuv backbone, forward primer |
| UctC_R | ATCCTTACTCGAGTTTGGATCCTCACAGCACGCCCGAG | Amplify *uctC* from DSM 29578[2] gDNA for restriction digest cloning into pBb(RK2)1k-GFPuv backbone, reverse primer |
| HutH2_F | AATTTCAGAATTCAAAAGATCTTTTAAGAAGGAGATATACATGTGACTGCGCTAAATCTG | Amplify *hutH2* from ATCC 9447 gDNA for restriction digest cloning into pBb(RK2)1k-GFPuv backbone, forward primer |
| HutH2_R | ATCCTTACTCGAGTTTGGATCCTTACAGGCTCGGCAGC | Amplify *hutH2* from ATCC 9447 gDNA for restriction digest cloning into pBb(RK2)1k-GFPuv backbone, reverse primer |
| pBb(RK2)1k_F | GGATCCAAACTCGAGTAAG | Amplify pBb(RK2)1k-GFPuv backbone for HiFi assembly, forward primer |
| pBb(RK2)1k_R | ATGTATATCTCCTTCTTAAAAGATCT | Amplify pBb(RK2)1k-GFPuv backbone for HiFi assembly, reverse primer |
| YbhH-HiFi_F | TTTAAGAAGGAGATATACATATGTCTTTTGAACTGGACCTTCCC | Amplify *ybhH* from DSM 21509 gDNA for HiFi assembly, forward primer |
| YbhH-HiFi_R | CCTTACTCGAGTTTGGATCCTTAGCCCCGCCCTTTCAAC | Amplify *ybhH* from DSM 21509 gDNA for HiFi assembly, reverse primer |
| Seq-pBb(Rk2)1k_F | CAATTAATCATCCGGCTCG | Forward sequencing primer for overexpression plasmids |
| Seq-pBb(Rk2)1k_R | GACTCTAGTAGAGAGCGTTC | Reverse sequencing primer for overexpression plasmids |

**Table S5. List of plasmids used in this study.**

| Plasmid name | Description | Source |
|---|---|---|
| pBb(RK2)1k-GFPuv | IPTG-inducible trc promoter expressing *gfpuv*, kanamycin resistance, RK2 origin of replication | Cook et al. 2018 |
| pBb(RK2)1k-PS__04252 | IPTG-inducible trc promoter expressing PS__04252 amplified from DSM 21509, kanamycin resistance, RK2 origin of replication | This study |
| pBb(RK2)1k-PS__04251 | IPTG-inducible trc promoter expressing PS__04251 amplified from DSM 21509, kanamycin resistance, RK2 origin of replication | This study |
| pBb(RK2)1k-YbhH | IPTG-inducible trc promoter expressing *ybhH* amplified from DSM 21509, kanamycin resistance, RK2 origin of replication | This study |
| pBb(RK2)1k-RhtA | IPTG-inducible trc promoter expressing *rhtA* amplified from DSM 29578[2], kanamycin resistance, RK2 origin of replication | This study |
| pBb(RK2)1k-UctC | IPTG-inducible trc promoter expressing *uctC* from amplified from DSM 29578[2], kanamycin resistance, RK2 origin of replication | This study |
| pBb(RK2)1k-HutH2 | IPTG-inducible trc promoter expressing *hutH2* amplified from ATCC 9447, kanamycin resistance, RK2 origin of replication | This study |
| pBb(RK2)1k-ProY1 | IPTG-inducible trc promoter expressing *proY_1* amplified from ATCC 9447, kanamycin resistance, RK2 origin of replication | This study |
