## Supplementary File 6 for "Population genomics-guided engineering of phenazine biosynthesis in *Pseudomonas chlororaphis*": phenotype_collapsing.py

```
# -*- coding: utf-8 -*-
"""
@author: Varun Trivedi
"""
# Refining the preliminary list of significant hits to create a 'phenotype-collapsed' list

import csv

tab = 'preliminary_list_of_hits.tab' # .tab file containing the preliminary list of hits
# Lists for storing various unitig attributes
seq = []; effect = []; length = []; q = []; chrom = []; start = []; end = [];
region = []; phen = []

with open(tab,'r') as f:
    reader = csv.reader(f, delimiter = '\t')
    for row in reader: # Reading contents of the .tab file
        seq.append(row[0]); effect.append(row[1])
        q.append(row[2]); length.append(row[3])
        chrom.append(row[4])
        start.append(row[5]); end.append(row[6])
        region.append(row[7]); phen.append(row[8])
# Excluding column titles from the respective lists
seq.remove(seq[0]); effect.remove(effect[0]); q.remove(q[0]);
length.remove(length[0]); chrom.remove(chrom[0]); start.remove(start[0]);
end.remove(end[0]); region.remove(region[0]); phen.remove(phen[0]);

hits = []; # List storing attributes of each hit in the preliminary list
# Each element of hits[] corresponds to all attributes of a single hit entry
# For each such element, all attributes are concatenated into a single string separated by '+'
for i in range(len(chrom)):
	hits.append(str(length[i])+'+'+chrom[i]+'+'+str(start[i])+'+'+str(end[i])+'+'+region[i]+'+'+seq[i]+'++'+phen[i]+'+'+str(effect[i])+'+'+str(q[i]))
short_hits = []; # List storing a truncated version of each element in hits[]
# Each element of short_hits[] would thus consist of all hit attributes except for phenotype, effect & q-value
# (as these would have different values for the same k-mer/unitig across different phenotypes)

# Unitigs/k-mers found in different phenotype lists that have the same length and sequence
# & map to the same chromosome & region (CDS/non-coding) are collapsed into a single entry (i.e., a unique hit)
for h in hits:
	short_hits.append(h[:h.find('++')])
unique_hits = list(set(short_hits)); # List that stores each unique hit entry for the phenotype-collapsed list
count = [];
# Each unique hit entry is assigned a score based on the number of phenotype lists the hit (unitig/k-mer) is found (to be significant) in.
# This score is called the phenotype score. count[] stores this phenotype score for each hit.
length_2 = []; chrom_2 = []; start_2 = []; end_2 = []; region_2 = []; seq_2 = []
phen_2 = []; q_2 = []; effect_2 = []; sd_effect = []
# The strings consisting of concatenated attributes (separated by '+') are fragmented & individual attributes
# are stored in the lists above for each hit
for i in unique_hits:
	c = 0 # Each hit is initially assigned a phenotype score of 0
	indices = [j for j, x in enumerate(i) if x == '+']
	length_2.append(i[:indices[0]]); chrom_2.append(i[indices[0]+1:indices[1]])
	start_2.append(i[indices[1]+1:indices[2]]); end_2.append(i[indices[2]+1:indices[3]])
	region_2.append(i[indices[3]+1:indices[4]]); seq_2.append(i[indices[4]+1:])
	phenotype = ''; eff = []; q_val = []
	for h in hits:
		if i in h:
			y = h[h.find('++')+2:]
			indices_2 = [j for j, x in enumerate(y) if x == '+']
			phenotype += y[:indices_2[0]]+', ' # 'phenotype' stores names of all phenotypes a hit is found (significant) in
			eff.append(float(y[indices_2[0]+1:indices_2[1]]))
			q_val.append(float(y[indices_2[1]+1:]))
			c += 1 # Incrementing phenotype score by 1 every time a hit (unitig/k-mer) is found in the preliminary list
	count.append(c); phen_2.append(phenotype[:-2]); 
# Effect of each unique hit = 
#                   Max. of effects of the hit in all phenotype lists (if effect > 0)
#                   Min. of effects of the hit in all phenotype lists (if effect < 0)
	if eff[0] > 0: effect_2.append(max(eff))
	else: effect_2.append(min(eff))
# q-value of each unique hit = Min. of q-values of the hit in all phenotype lists
	q_2.append(min(q_val))

g = 'phenotype_collapsed_list.tab'
with open(g,'w', newline = '') as f: # Writing results in a new .tab file
    writer = csv.writer(f, delimiter = '\t')
    writer.writerow(['chromosome','start','end','region','phenotype_score','phenotype','effect','q-value','length','sequence'])
    for i in range(len(unique_hits)):
        row = []
        row.append(chrom_2[i]); row.append(start_2[i])
        row.append(end_2[i]); row.append(region_2[i])
        row.append(count[i]); row.append(phen_2[i])
        row.append(effect_2[i]); row.append(q_2[i])
        row.append(length_2[i]); row.append(seq_2[i])
        writer.writerow(row)
```
