## Supplementary File 6 for "Population genomics-guided engineering of phenazine biosynthesis in *Pseudomonas chlororaphis*": rev_comp_collapsing.py

```
# -*- coding: utf-8 -*-
"""
@author: Varun Trivedi
"""
# Refining the phenotype- + strain-collapsed list to create the final hit list
import csv

def rev_comp(a):
	""" Function to convert a unitig into its reverse complement """
	a = ''.join(reversed(a)) # reversing the unitig sequence
	x = {'A':'T','T':'A','C':'G','G':'C'}
	b = ''
	for i in range(len(a)):
		b += x[a[i]]
	return b

tab = 'phen_strain_collapsed_list.tab' # .tab file containing the phenotype- + strain-collapsed list
# Lists for storing various unitig attributes
seq = []; effect = []; length = []; q = []; chrom = []; start = []; end = [];
region = []; phen = []; score = []

with open(tab,'r') as f:
    reader = csv.reader(f, delimiter = '\t')
    for row in reader: # Reading contents of the .tab file
        chrom.append(row[0]); region.append(row[3])
        start.append(row[1]); end.append(row[2])
        score.append(row[4]); phen.append(row[5])
        effect.append(row[6]); q.append(row[7])
        length.append(row[8]); seq.append(row[9])
# Excluding column titles from the respective lists
seq.remove(seq[0]); effect.remove(effect[0]); q.remove(q[0]);
length.remove(length[0]); chrom.remove(chrom[0]); start.remove(start[0]);
end.remove(end[0]); region.remove(region[0]); phen.remove(phen[0]); score.remove(score[0])

unitig_features = []; # List for storing unitig attributes like length, effect, score, final score, sequence & region
for i in range(len(region)):
	unitig_features.append(region[i]+'+'+str(score[i])+'+'+phen[i]+'+'+str(effect[i])+'+'+str(q[i])+'+'+str(length[i])+'+'+str(score[i])+'+'+seq[i])
for i in range(len(unitig_features)):
# If, for a unitig in 'unitig_features[]', a second unitig with identical attributes & reverse-complemented sequence is found in the same list,
# the string at the position of the second unitig in the list is replaced with '0' to indicate that it should be excluded from the final list
	if unitig_features[i] != '0':
		x = unitig_features[i].rfind('+') # Finding the position of '+' appearing just before the unitig sequence
		if unitig_features[i][:x+1] + rev_comp(seq[i]) in unitig_features:
			y = unitig_features.index(unitig_features[i][:x+1] + rev_comp(seq[i])) # Position of the unitig with the reverse-complemented sequence, if it is found in the list
			unitig_features[y] = '0'

g = 'final_list.tab'
with open(g,'w', newline = '') as f: # Writing results in a new .tab file
    writer = csv.writer(f, delimiter = '\t')
    writer.writerow(['chromosome','start','end','region','phenotype_score','phenotype','effect','q-value','length','sequence'])
    #c = 0
    for i in range(len(unitig_features)):
        if unitig_features[i] != '0':
            row = []
            row.append(chrom[i]); row.append(start[i])
            row.append(end[i]); row.append(region[i])
            row.append(score[i]); row.append(phen[i])
            row.append(effect[i]); row.append(q[i])
            row.append(length[i]); row.append(seq[i])
            writer.writerow(row)
```
