## Supplementary File 6 for "Population genomics-guided engineering of phenazine biosynthesis in *Pseudomonas chlororaphis*": strain_collapsing.py

```
# -*- coding: utf-8 -*-
"""
@author: Varun Trivedi
"""
# Refining the phenotype-collapsed list to create a phenotype- + strain-collapsed list

import csv

tab = 'phenotype_collapsed_list.tab' # .tab file containing the phenotype-collapsed list
# Lists for storing various unitig attributes
seq = []; effect = []; length = []; q = []; chrom = []; start = []; end = [];
region = []; phen = []; score = []

with open(tab,'r') as f:
    reader = csv.reader(f, delimiter = '\t')
    for row in reader: # Reading contents of the .tab file
        chrom.append(row[0]); region.append(row[3])
        start.append(row[1]); end.append(row[2])
        score.append(row[4]); phen.append(row[5])
        effect.append(row[6]); q.append(row[7])
        length.append(row[8]); seq.append(row[9])
# Excluding column titles from the respective lists
seq.remove(seq[0]); effect.remove(effect[0]); q.remove(q[0]);
length.remove(length[0]); chrom.remove(chrom[0]); start.remove(start[0]);
end.remove(end[0]); region.remove(region[0]); phen.remove(phen[0]); score.remove(score[0])

hits = []; # List storing attributes of each hit in the processed list
# Each element of hits[] corresponds to all attributes of a single hit entry
# For each such element, all attributes are concatenated into a single string separated by '+'
for i in range(len(chrom)):
	hits.append(chrom[i]+'+'+str(start[i])+'+'+str(end[i])+'++'+region[i]+'+'+str(score[i])+'+'+phen[i]+'+'+str(effect[i])+'+'+str(q[i])+'+'+str(length[i])+'+'+seq[i])
short_hits = []; # List storing a truncated version of each element in hits[]
# Each element of short_hits[] would thus consist of all hit attributes except for chromosome, start & end position
# (as these would have different values for the same k-mer/unitig across different strains in which that unitig/k-mer is found)

# Unitigs mapping to the same CDS, having the same phen. score, length, sequence, avg. effect, & q-value
# and found in the same set of phenotypes are collapsed into a single entry (i.e., a unique hit)
for h in hits:
	short_hits.append(h[h.find('++')+2:])
unique_hits = list(set(short_hits)); # List that stores each unique hit entry for the phenotype- + strain-collapsed list
length_2 = []; chrom_2 = []; start_2 = []; end_2 = []; region_2 = [];
phen_2 = []; q_2 = []; effect_2 = []; score_2 = []; seq_2 = []
# The strings consisting of concatenated attributes (separated by '+') are fragmented & individual attributes
# are stored in the lists above for each hit
for i in unique_hits:
	indices = [j for j, x in enumerate(i) if x == '+']
	region_2.append(i[:indices[0]]); score_2.append(i[indices[0]+1:indices[1]])
	phen_2.append(i[indices[1]+1:indices[2]]); effect_2.append(i[indices[2]+1:indices[3]])
	q_2.append(i[indices[3]+1:indices[4]]); 
	length_2.append(i[indices[4]+1:indices[5]]); seq_2.append(i[indices[5]+1:])
	for h in hits:
		if i in h:
			y = h[:h.find('++')]
			indices_2 = [j for j, x in enumerate(y) if x == '+']
			chrom_2.append(y[:indices_2[0]])
			start_2.append(y[indices_2[0]+1:indices_2[1]])
			end_2.append(y[indices_2[1]+1:])
			break

g = 'phen_strain_collapsed_list.tab'
with open(g,'w', newline = '') as f: # Writing results in a new .tab file
    writer = csv.writer(f, delimiter = '\t')
    writer.writerow(['chromosome','start','end','region','phenotype_score','phenotype','effect','q-value','length','sequence'])
    for i in range(len(unique_hits)):
        row = []
        row.append(chrom_2[i]); row.append(start_2[i])
        row.append(end_2[i]); row.append(region_2[i])
        row.append(score_2[i]); row.append(phen_2[i])
        row.append(effect_2[i]); row.append(q_2[i])
        row.append(length_2[i]); row.append(seq_2[i])
        writer.writerow(row)
```
